# The MEK-ERK signaling pathway promotes maintenance of cardiac chamber identity

**DOI:** 10.1101/2023.07.17.549343

**Authors:** Yao Yao, Deborah Yelon

## Abstract

Ventricular and atrial cardiac chambers have unique structural and contractile characteristics that underlie their distinct functions. Intriguingly, the maintenance of chamber-specific features requires active reinforcement, even in differentiated cardiomyocytes. Prior studies in zebrafish have shown that sustained FGF signaling acts upstream of *nkx2.5* to maintain ventricular identity, but the rest of this maintenance pathway remains unclear. Here, we show that MEK1/2-ERK1/2 signaling acts downstream of FGF and upstream of *nkx2.5* to promote ventricular maintenance. Inhibition of MEK signaling, like inhibition of FGF signaling, results in ectopic atrial gene expression and reduced ventricular gene expression in ventricular cardiomyocytes. FGF and MEK signaling both influence ventricular maintenance over a similar timeframe, when phosphorylated ERK (pERK) is present in the myocardium. However, the role of FGF-MEK activity seems to be context-dependent: some ventricular regions are more sensitive than others to inhibition of FGF-MEK signaling. Additionally, in the atrium, although endogenous pERK does not induce ventricular traits, heightened MEK signaling can provoke ectopic ventricular gene expression. Together, our data reveal chamber-specific roles of MEK-ERK signaling in the maintenance of ventricular and atrial identities.

**SUMMARY STATEMENT:** The MEK-ERK signaling pathway plays distinct roles in the maintenance of ventricular and atrial cardiomyocyte identities.

## INTRODUCTION

The vertebrate heart contains two types of cardiac chambers with distinct functions: atrial chambers collect and supply blood to the ventricles, whereas ventricular chambers forcefully propel blood to the periphery. The morphological, physiological, and molecular differences between the ventricular and atrial myocardium – including differences in tissue architecture, conductive properties, and gene expression profiles – underlie the functional characteristics of each chamber (Asp et al., 2012; DeLaughter et al., 2016; Li et al., 2016; Moorman and Christoffels, 2003; Singh et al., 2016; van Weerd and Christoffels, 2016; Wessels and Sedmera, 2004). Although the operational importance of chamber-specific traits is well established, we do not yet fully understand how ventricular and atrial cardiomyocytes acquire and maintain their characteristic features.

Fate mapping studies in chick, zebrafish, and mouse have suggested that chamber fate specification occurs in the early embryo, where spatial segregation of ventricular and atrial myocardial lineages is evident prior to or during gastrulation (Bardot et al., 2017; Garcia-Martinez and Schoenwolf, 1993; Ivanovitch et al., 2021; Keegan et al., 2004; Rosenquist, 1970; Stainier et al., 1993). During these stages, the signaling pathways that pattern the embryonic axes, including Nodal, FGF, and BMP signaling, also seem to influence patterning of the cardiac progenitors (Keegan et al., 2004; Marques and Yelon, 2009; Marques et al., 2008; Reiter et al., 2001). Interestingly, despite early specification of chamber progenitors, the chamber identities of differentiated cardiomyocytes require active maintenance as development proceeds (Martin and Waxman, 2021; Yao et al., 2021). For example, several transcription factors have been shown to be required for maintenance of chamber-specific gene expression patterns. In mouse, *COUP-TFII* and *TBX5* are essential for atrial identity maintenance: both factors sustain the expression of atrial genes and repress the expression of ventricular genes in differentiated atrial cardiomyocytes (Sweat et al., 2023; Wu et al., 2013). Likewise, in zebrafish, *Nr2f1a*, a functional homolog of mammalian *COUP-TFII*, prevents atrial cardiomyocytes from developing ectopic ventricular and pacemaker cell identities (Martin et al., 2023). Conversely, studies in mouse and chick indicate critical roles of *Hey2 and Irx4* in ventricular identity maintenance, as they repress expression of atrial genes in the ventricular myocardium (Bao et al., 1999; Bruneau et al., 2001; Koibuchi and Chin, 2007; Xin et al., 2007). While these transcription factors clearly promote maintenance of chamber identities, less is known about the regulatory pathways that control their chamber-specific expression.

It is therefore intriguing that prior studies have indicated a role for FGF signaling in promoting the expression of Nkx transcription factors, which serve as upstream regulators of *Hey2* and *Irx4*. In zebrafish, mouse, and hESC models, *Nkx2-5* homologs are required to drive expression of *Hey2* and *Irx4* homologs in the ventricle (Anderson et al., 2018; Bruneau et al., 2000; Targoff et al., 2013). Furthermore, in zebrafish, loss of function of the Nkx factors *nkx2.5* and *nkx2.7* disrupts ventricular identity maintenance, causing the acquisition of expression of the atrial gene *amhc* (*atrial myosin heavy chain,* also known as *myh6*) in the ventricle, coupled with the loss of expression of the ventricular gene *vmhc* (*ventricular myosin heavy chain,* also known as *myh7*) (Targoff et al., 2013). Similarly, inhibition of FGF signaling after the initiation of myocardial differentiation results in ectopic expression of *amhc* and reduced expression of *vmhc* in the ventricle, as well as reduced expression of *nkx2.5* and *nkx2.7* (Pradhan et al., 2017). Since overexpression of *nkx2.5* can partially rescue the ventricular defects caused by inhibition of FGF signaling (Pradhan et al., 2017), the synthesis of these data demonstrates that FGF signaling plays an important role, upstream of Nkx factors, in maintaining ventricular chamber identity. However, it remains unclear how FGF signaling regulates the expression of Nkx factors and whether additional genes also mediate the role of FGF signaling in reinforcement of ventricular traits.

In many developmental processes, FGF signaling is known to execute its functions by activating the MEK1/2-ERK1/2 pathway (Brewer et al., 2016; Dailey et al., 2005). For instance, FGF4 functions through the ERK1/2 pathway to activate primitive endoderm specification within the inner cell mass of the mouse embryo (Kang et al., 2013; Krawchuk et al., 2013; Nichols et al., 2009; Soszyńska et al., 2019), where live visualization of ERK activity has revealed distinct ERK signaling dynamics in different lineages (Simon et al., 2020). As another example, studies in multiple vertebrate models have indicated a role of ERK1/2 downstream of FGF signaling in somite segmentation (Akiyama et al., 2014; Delfini et al., 2005; Niwa et al., 2011). In contrast, FGF signaling in other developmental contexts is mediated by different signal transduction mechanisms including the PI3K-AKT/PKB and PLCγ-PKC pathways, as well as pathways using MAPK proteins other than ERK1/2 (Brewer et al., 2016; Dailey et al., 2005). For example, in *Xenopus*, FGF signaling acts via p38 MAPK to regulate the initial expression of *nkx2.5* in cardiac progenitors (Keren-Politansky et al., 2009). FGF signaling has also been shown to function through PI3K pathways in the development of GnRH-secreting neurons and through JNK pathways during bile acid biosynthesis (Brewer et al., 2016; Holt et al., 2003; Hu et al., 2013). It therefore seems likely that one of these established downstream signal transduction pathways facilitates the role of FGF signaling in the maintenance of ventricular identity.

Here, we modulate the MEK1/2-ERK1/2 signaling pathway, using both pharmacological and genetic approaches, and evaluate the impact on chamber identity maintenance. We find that inhibition of MEK-ERK signaling following myocardial differentiation results in ectopic *amhc* expression as well as reduced *vmhc* expression in ventricular cells, similar to the effects of inhibition of FGF signaling. Additionally, MEK-ERK signaling and FGF signaling appear to be required for ventricular identity maintenance during similar timeframes, correlating with stages when phosphorylated ERK is detectable in the myocardium. Epistasis analysis indicates that a constitutively active form of MEK1 can partially rescue the effects of FGF pathway inhibition and that overexpression of *nkx2.5* can partially rescue the effects of MEK pathway inhibition. Together, these data support a model in which MEK-ERK signaling acts downstream of FGF signaling and upstream of *nkx* factors to promote ventricular identity maintenance. Intriguingly, our results also indicate that the role of this FGF-MEK-ERK-*nkx* pathway is context-dependent: some regions of the ventricle are more sensitive than other regions to inhibition of FGF or MEK signaling. Additionally, in the atrium, although endogenous levels of MEK-ERK signaling are not sufficient to induce ventricular traits, heightened MEK signaling can cause ectopic ventricular gene expression. Thus, our studies indicate distinct functions of MEK-ERK signaling in ventricular and atrial identity maintenance.

## RESULTS

### MEK signaling, like FGF signaling, promotes ventricular identity maintenance

To investigate the pathways that might mediate FGF signal transduction during ventricular identity maintenance, we began by evaluating the effects of a set of kinase inhibitors on gene expression patterns in the ventricle. We found that inhibition of PI3K or JNK, using LY294002 or SP600125, failed to induce any ectopic *amhc* in the ventricle (Fig. S1). In contrast, treatment with PD0325901, an inhibitor of MEK1/2 activity that prevents the phosphorylation of ERK1/2 (Sebolt-Leopold, 2008), induced both ectopic *amhc* transcripts (Fig. 1B) and Amhc protein (Fig. 1E) in the ventricle. Moreover, the pattern of ectopic atrial marker localization in PD0325901-treated embryos was highly reminiscent of the phenotype that we had previously observed in embryos treated with the FGFR inhibitor SU5402 (Fig. 1C,F) (Pradhan et al., 2017). Additionally, we found that MEK pathway inhibition reduced the extent and intensity of *vmhc* expression in the ventricle (Fig. 1H), similar to the effects of FGF pathway inhibition with SU5402 (Fig. 1I) or with expression of a dominant-negative form of FGFR1 (Pradhan et al., 2017). Together, these data suggest that MEK-ERK signaling, like FGF signaling, acts both to prevent atrial gene expression and to promote ventricular gene expression within the ventricle.

**Figure 1.**
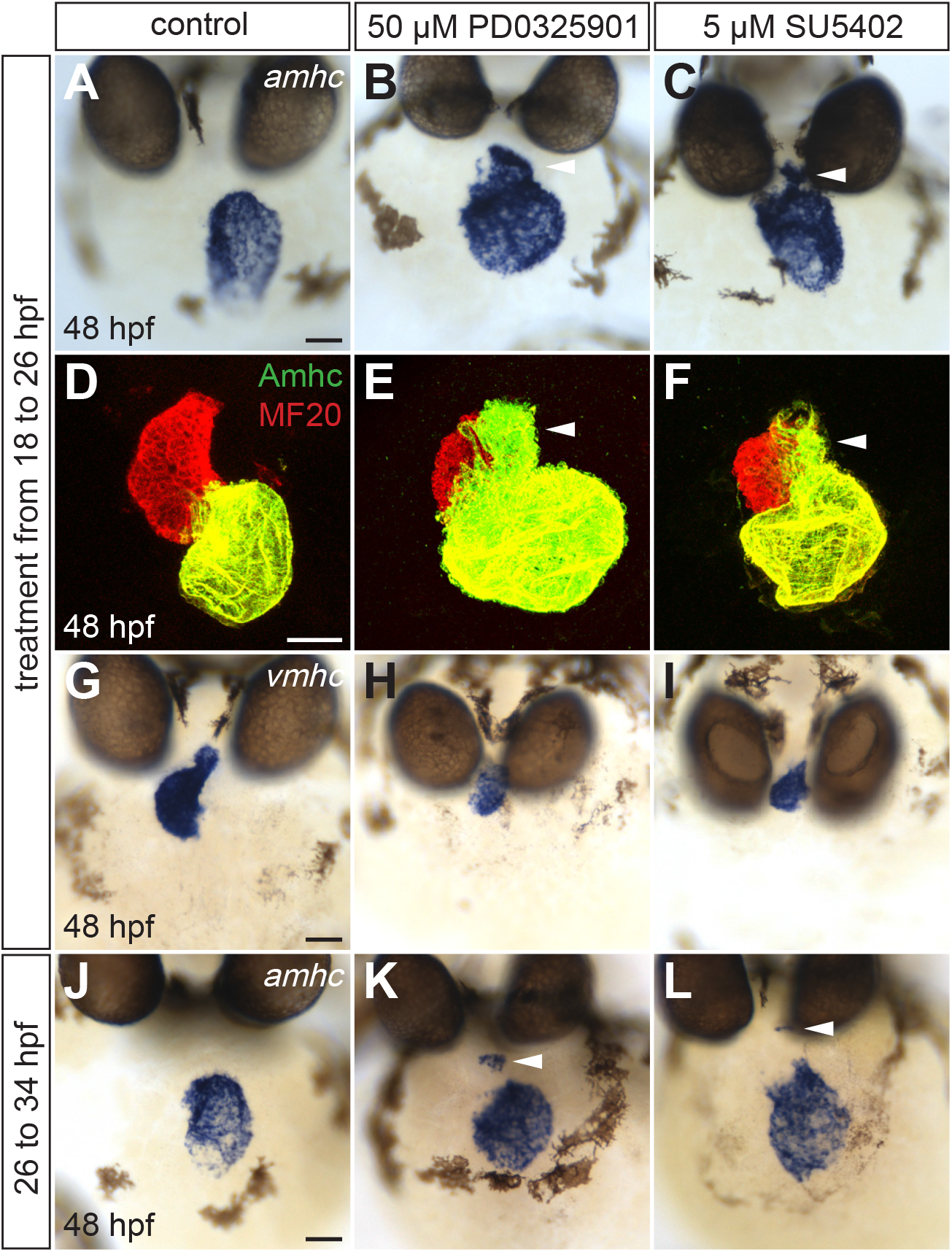
MEK signaling promotes ventricular identity maintenance. *In situ* hybridization shows expression of *amhc* (A-C, J-L) or *vmhc* (G-I), and immunofluorescence (D-F) shows distribution of Amhc (green) within the myocardium (labeled with MF20, red), in frontal views of the heart at 48 hpf. Wild-type embryos were treated with DMSO, PD0325901 or SU5402 from 18 to 26 hpf (A-I) or from 26 to 34 hpf (J-L). In PD0325901-treated (B,E,K) and SU5402-treated embryos (C,F,L), ectopic *amhc* transcripts or Amhc protein appear in similar locations (arrowheads) within the ventricle. Additionally, PD0325901-treated (H) and SU5402-treated (I) embryos display reduced *vmhc* expression, compared to DMSO-treated controls (G). (A,B,D,F,K,L) n>13, (C) n=7, (E) n=6, (G,J) n=10, (H) n=12, (I) n=13. Scale bars: 50 μm.

Extending our comparison of the influences of MEK and FGF signaling on the ventricular myocardium, we examined whether these pathways act during the same time interval. Treatment with PD0325901 or SU5402 starting at 18 hours post fertilization (hpf), when ventricular and atrial myocardial differentiation are already underway (Berdougo et al., 2003; Yelon et al., 1999), effectively induced ectopic *amhc* expression in the ventricle (Fig. 1A-F). However, treatment at later time points, such as 26 hpf, was less potent, causing ectopic *amhc* expression only in a few cells, primarily near the outflow tract (OFT) (Fig. 1K,L). Thus, our data suggest that MEK and FGF signaling are both required during a comparable timeframe to promote ventricular identity maintenance.

Interestingly, not all of the regions in the ventricle seemed equally sensitive to the inhibition of MEK or FGF signaling. Typically, we noted that cells in the ventricular inner curvature (IC) and OFT more readily expressed ectopic *amhc*, while *amhc* expression was less frequently observed in the ventricular outer curvature (OC) (Fig. 1B,C,E,F). This general trend was consistent with our prior observations of the effects of FGF pathway inhibition (Pradhan et al., 2017). Investigating this differential sensitivity more systematically, we found that low concentrations of SU5402 or PD0325901 frequently induced ectopic *amhc* expression in the OFT and in the ventricular apex (Fig. 2B,G). Higher concentrations led to the additional induction of ectopic *amhc* in the IC (Fig. 2C,H) and the central portion of the ventricle (Fig. 2D,I). With the highest concentrations tested, ectopic *amhc* was also observed in the OC (Fig. 2E,J), such that all regions of the ventricle were expressing *amhc*. Correspondingly, we observed greater loss of *vmhc* expression when treating embryos with higher concentrations of SU5402 or PD0325901 (Fig. 2N-Q,S-V).

**Figure 2.**
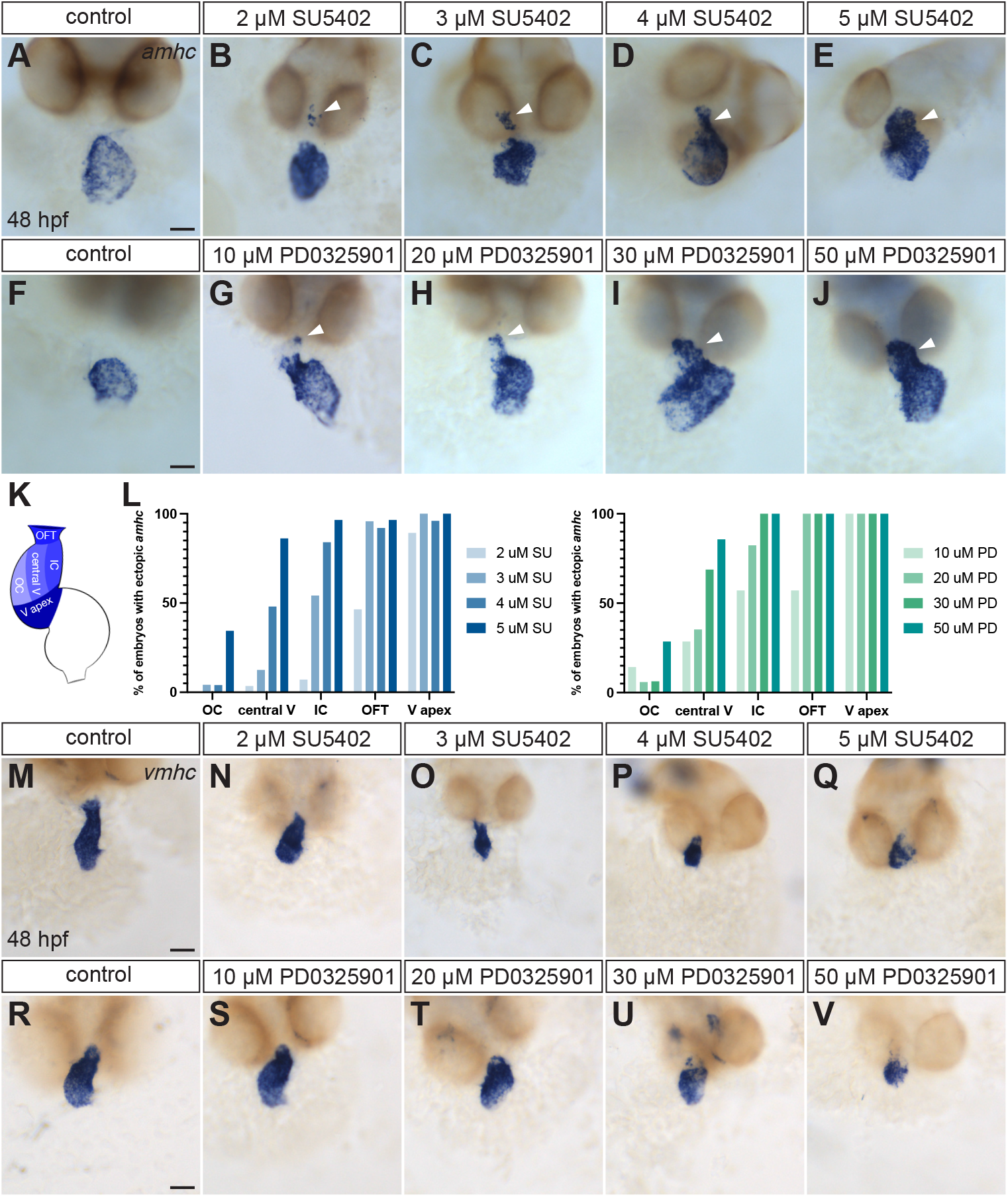
Ventricular regions are differentially sensitive to the inhibition of FGF signaling or MEK signaling. Expression of *amhc* (A-J) or *vmhc* (M-V) at 48 hpf (as in Fig. 1) shows effects of treatment of wild-type embryos with a range of concentrations of SU5402 or PD0325901 from 18 to 26 hpf. Representative examples demonstrate that higher concentrations of either compound lead to broader ectopic expression of *amhc* (A-J, arrowheads), together with reduced expression of *vmhc* (M-V). (K) Diagram depicts locations of ventricular regions, with darker shading in regions that more readily express ectopic *amhc* when treated with SU5402 or PD0325901. Figure S2 provides examples of hearts with ectopic *amhc* in each of these regions. (L) Bar graphs show the percentage of embryos receiving each treatment that expressed ectopic *amhc* in each ventricular region. (A) n=21, (B) n=28, (C) n=24, (D) n=25, (E) n=29, (F,J) n=14, (G) n=7, (H) n=17, (I) n=16, (M) n=8, (N,Q,R) n=12, (O) n=6, (P,T,U) n=13, (S,V) n=15. Scale bars: 50 μm.

Altogether, for a given concentration of either SU5402 or PD0325901, more embryos expressed ectopic *amhc* in the more sensitive regions, such as the OFT, while fewer embryos expressed *amhc* in the less sensitive regions, such as the OC (Fig. 2L). Additionally, higher concentrations of either inhibitor generally resulted in more embryos expressing *amhc* in any ventricular region (Fig. 2L). These dose-dependent responses suggest differential degrees of plasticity for ventricular identity in different portions of the ventricle. Moreover, the phenotypic similarities between SU5402-treated and PD0325901-treated ventricles suggest that FGF and MEK signaling could act in the same pathway to regulate chamber-specific characteristics.

### MEK-ERK signaling is active throughout the myocardium

Our prior work has suggested that FGF signaling is required cell-autonomously within ventricular cardiomyocytes to maintain their identity (Pradhan et al., 2017). Ectopic *amhc* expression in the *fgf8a* mutant ventricle suggests that Fgf8a is one of the critical signals involved (Pradhan et al., 2017), and *fgf8a* is expressed in the myocardium during the 18-26 hpf time interval that seems to be critical for ventricular identity maintenance (Reifers et al., 2000). We therefore hypothesized that FGF signaling activates MEK-ERK signaling within the ventricular myocardium during these stages. In order to visualize MEK-ERK signaling activity, we went on to examine the localization of phosphorylated ERK1/2 (pERK) in the cardiac cone at 20 hpf and in the heart tube at 24 hpf.

Heart tube assembly begins with the fusion of bilateral populations of myocardial precursors to form the cardiac cone at the embryonic midline, with ventricular cardiomyocytes at the center of this shallow cone, and atrial cardiomyocytes at the periphery (Berdougo et al., 2003; Staudt and Stainier, 2012). In wild-type embryos, we detected pERK localization in multiple cells within the myocardial cone, as well as within its endocardial core (Fig. 3A,B). This pERK signal was dramatically diminished in PD0325901-treated embryos (Fig. S3), validating the specificity of our anti-pERK immunofluorescence protocol. Notably, the number and location of pERK+ cardiomyocytes and the intensity of the pERK signal varied widely between samples (Figs 3B,E, S4). We did not identify any consistent regional patterns of pERK localization within the myocardium: there was no consistently detectable enrichment of pERK levels in the more central ventricular cells or in the more peripheral atrial cells, nor was there any consistent enhancement of pERK signal on the left or right sides or in the anterior or posterior portions of the cardiac cone (Fig. S4). Altogether, the localization of pERK appeared unique in each embryo. Given the broad variability of pERK distribution, we suspect that MEK-ERK signaling is highly dynamic within the myocardium of the cardiac cone.

**Figure 3.**
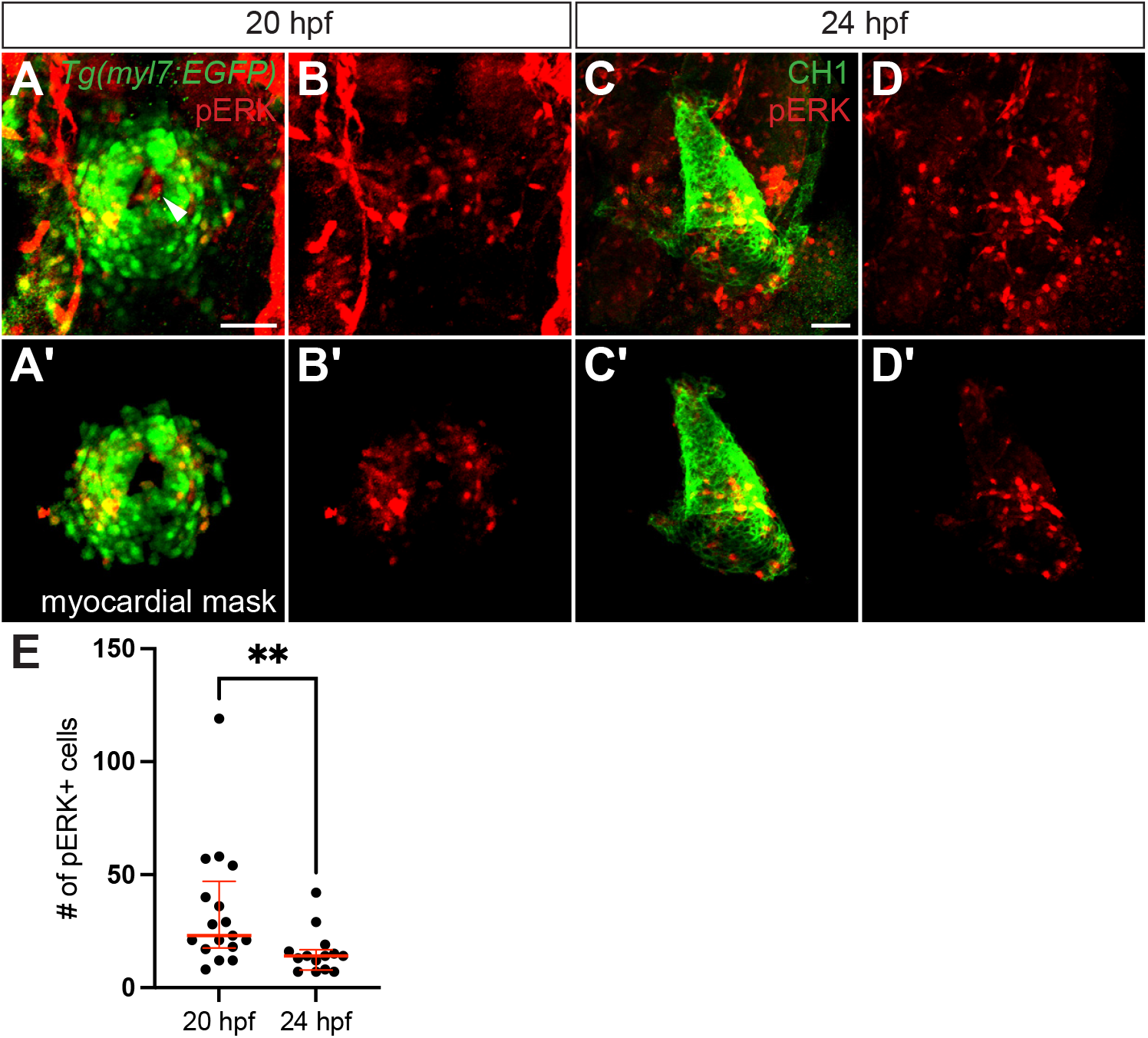
MEK-ERK signaling activity is detectable in the myocardium. Three-dimensional reconstructions show pERK (red) localization within the myocardium, labeled with *Tg(myl7:EGFP)* (green) at 20 hpf (A,B) or CH1 (green; anti-tropomyosin) at 24 hpf (C,D). Dorsal views, anterior up (A,B) or arterial pole up (C,D). pERK is also detectable in other tissues, including the endocardium (arrowhead in A). (A’-D’) Myocardial masks (see Materials and Methods) include only the pERK labeling that overlaps with myocardial labeling. (A,B) n=17, (C,D) n=7. Scale bars: 50 μm. (E) Graph compares the numbers of pERK+ cardiomyocytes at 20 and 24 hpf. We often found fewer pERK+ cardiomyocytes at 24 hpf than at 20 hpf; this reduction likely reflects a more limited distribution of ERK signaling activity at 24 hpf, but we note that it could also result from differential sensitivity of the distinct laser power and exposure settings used when imaging embryos at these two stages. Data at 24 hpf include 7 samples labeled with CH1 and 7 samples labeled with *Tg(myl7:EGFP)*. Red lines represent median and interquartile range. **p=0.0044, two-tailed Mann-Whitney test.

We continued to observe pERK in the myocardium after heart tube assembly was complete (Fig. 3C-E). While myocardial pERK was present in all embryos examined at 24 hpf, endocardial pERK was less frequently observed at this stage (only in 6/14 embryos). Additionally, although we did detect pERK in both the ventricular and atrial portions of the heart tube, we typically observed enriched pERK localization in the atrial myocardium (Fig. 3C,D). Thus, ERK signaling seems to be more broadly distributed in the cardiac cone than in the heart tube, with ERK signaling in the ventricular myocardium more prevalent in the cone than in the tube, correlating with the timeframe when FGF and MEK signaling are required for ventricular maintenance. Interestingly, the presence of pERK in the atrial myocardium is evidently not sufficient to induce ventricular traits.

### MEK-ERK signaling functions downstream of FGF signaling to repress ectopic *amhc* expression in the ventricle

To test if MEK signaling acts downstream of FGF signaling during ventricular identity maintenance, we next examined whether constitutive MEK1 activity could rescue the effects of FGF pathway inhibition. For this purpose, we generated stable lines carrying the transgene *Tg(hsp:MEK1^S219D^-mCherry)* (hereafter referred to as *Tg(hsp:caMEK1)*), which expresses a constitutively active form of MEK1 (caMEK1) in response to heat shock. Ser219 is predicted to be a key phosphorylation site during activation of zebrafish MEK1, and the substitution of Ser to Asp to mimic phosphorylation has been shown to increase ERK phosphorylation (Bolcome and Chan, 2010; Chou et al., 2015). Correspondingly, we found that heat-induced expression of *Tg(hsp:caMEK1)* significantly increased the numbers of pERK+ cardiomyocytes and the levels of pERK signal in the myocardium, although the degree of increase varied (Fig. 4).

**Figure 4.**
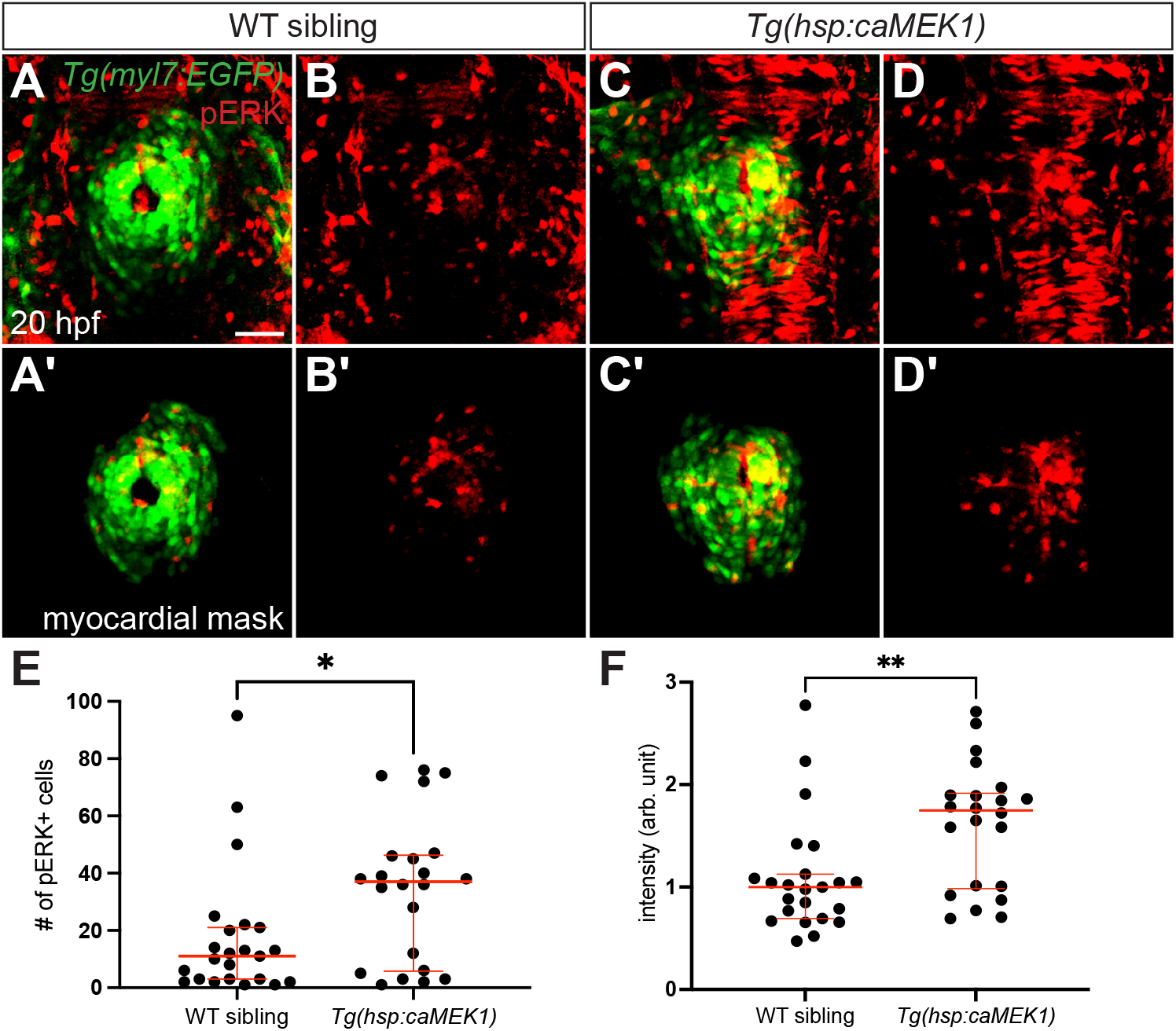
Constitutive activity of MEK1 increases pERK levels in the myocardium. Three-dimensional reconstructions compare pERK localization at 20 hpf (as in Fig. 3A,B) in *Tg(myl7:EGFP)* embryos (A,B) and *Tg(myl7:EGFP)* embryos carrying *Tg(hsp:caMEK1)* (C,D), after heat shock at 16 hpf. Myocardial masks (A’-D’) are used as in Fig. 3A’,B’. Expression of *Tg(hsp:caMEK1)* increases the amount of detectable pERK (C,D), compared to the endogenous level observed in wild-type (WT) sibling embryos (A,B). (A,B) n=24, (C,D) n=22. Scale bar: 50 μm. (E) Graph (as in Fig. 3E) compares the numbers of pERK+ cardiomyocytes in WT siblings and *Tg(hsp:caMEK1)* embryos. Note that the number of pERK+ cells in WT siblings here is lower than that in Fig. 3E. Because of the high intensity of pERK signal in *Tg(hsp:caMEK1)* embryos, we reduced our laser power and exposure settings in order to avoid saturating the confocal detector; the same imaging settings were used for WT siblings and *Tg(hsp:caMEK1)* embryos. (F) Graph compares the average intensity of myocardial pERK signal in each embryo examined. Each data point represents the mean of the intensity mean values for the pERK signal from all of the cardiomyocytes in a WT sibling or *Tg(hsp:caMEK1)* embryo; normalized data are expressed in arbitrary (arb.) units (see Materials and Methods). Red lines in graphs represent median and interquartile range. *p=0.0203, **p=0.0074; two-tailed Mann-Whitney test.

We evaluated whether expression of *Tg(hsp:caMEK1)* would prevent the appearance of ectopic *amhc* in the ventricular myocardium of SU5402-treated embryos. Indeed, following heat shock and SU5402 treatment, *Tg(hsp:caMEK1)* embryos displayed less ectopic *amhc* than their wild-type siblings (Fig. 5), suggesting that MEK-ERK signaling functions downstream of FGF signaling in the context of ventricular maintenance. When interpreting these results, we considered that SU5402 has also been shown to cause mild inhibition of VEGF signaling (Molina et al., 2007) and that the MEK-ERK pathway can also be activated by VEGF (Shin et al., 2016). However, treatment with a VEGFR inhibitor, SU4312, did not affect ventricular identity maintenance (Fig. S5), suggesting that our epistasis analysis reflects the importance of a FGF-MEK-ERK signaling pathway in repressing ectopic *amhc* expression in the ventricle.

**Figure 5.**
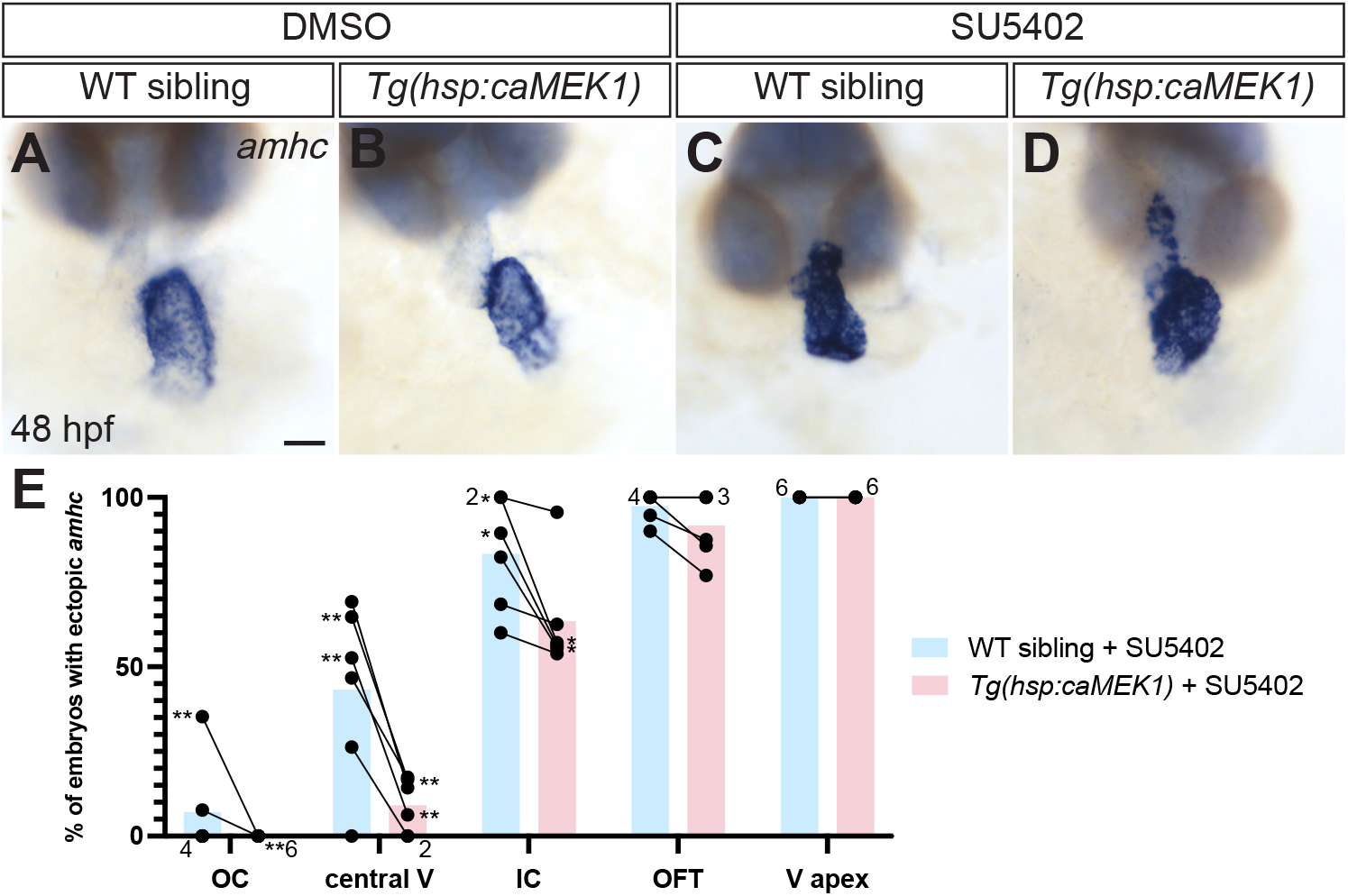
MEK-ERK signaling functions downstream of FGF signaling to maintain ventricular identity. (A-D) Expression of *amhc* at 48 hpf (as in Fig. 1) indicates effects of heat shock at 16 hpf followed by treatment with either DMSO (A,B) or 4-5 μM SU5402 (C,D) from 18 to 22 hpf. In SU5402-treated embryos, the ventricular distribution of ectopic *amhc* is less broad in embryos expressing *Tg(hsp:caMEK1)* (D) than in WT siblings (C). (E) Graph shows percentage of SU5402-treated embryos that expressed ectopic *amhc* in each ventricular region. Bars represent the mean values of the percentages of embryos expressing ectopic *amhc* from each of the six independent experiments that were performed. Lines connect the observations in *Tg(hsp:caMEK1)* embryos with the WT sibling controls in each replicate, and numerals next to data points indicate the number of replicates with the same value. Asterisks near the data points of a replicate pair indicate statistically significant differences between the expression of *amhc* in *Tg(hsp:caMEK1)* and WT sibling embryos. *p<0.05, **p<0.01; Fisher’s exact test. (A) n=18, (B) n=22, (C,D) n(WT sibling, *Tg(hsp:caMEK1)*)=(17,18), (13,7), (15,23), (19,16), (10,13), (19,8) in 6 independent experiments. Scale bar: 50 μm.

Interestingly, we noted that expression of *Tg(hsp:caMEK1)* was most potent in rescuing the less plastic ventricular regions, including the OC and central ventricle, of SU5402-treated embryos, while very little or no rescue was observed in the more plastic regions, such as the OFT and the ventricular apex (Fig. 5E). This pattern of partial rescue suggests the possibility that lower levels of MEK-ERK signaling are required for identity maintenance in the less plastic regions of the ventricle, whereas higher levels of MEK-ERK signaling are required in the more plastic regions.

### MEK-ERK signaling functions upstream of *nkx2.5* to repress ectopic *amhc* expression in the ventricle

When considering effector genes that could function downstream of the FGF-MEK-ERK pathway to promote ventricular maintenance, we kept in mind our previous finding that FGF signaling acts upstream of Nkx factors during this process (Pradhan et al., 2017). Similar to the effects of FGF pathway inhibition (Pradhan et al., 2017), MEK pathway inhibition resulted in reduced expression of both *nkx2.5* and *nkx2.7* (Fig. 6A-D), suggesting that MEK-ERK signaling might act in between FGF signaling and Nkx factors in ventricular identity maintenance. To test this, we examined whether overexpression of *nkx2.5* could prevent ectopic *amhc* expression in PD0325901-treated embryos. Following heat shock and PD0325901 treatment, we found that expression of *Tg(hsp:nkx2.5)* led to less ectopic *amhc* than was seen in wild-type siblings (Fig. 6E-I). Additionally, overexpression of *nkx2.5* achieved greater rescue in the less plastic regions of the ventricle, such as the OC and the central ventricle, whereas the observed rescue in the OFT and the ventricular apex was negligible (Fig. 6I). Together with our other results, these data support a model in which different regions of the ventricle have differential degrees of requirement for activation of a FGF-MEK-ERK-*nkx2.5* pathway in order to promote ventricular identity maintenance.

**Figure 6.**
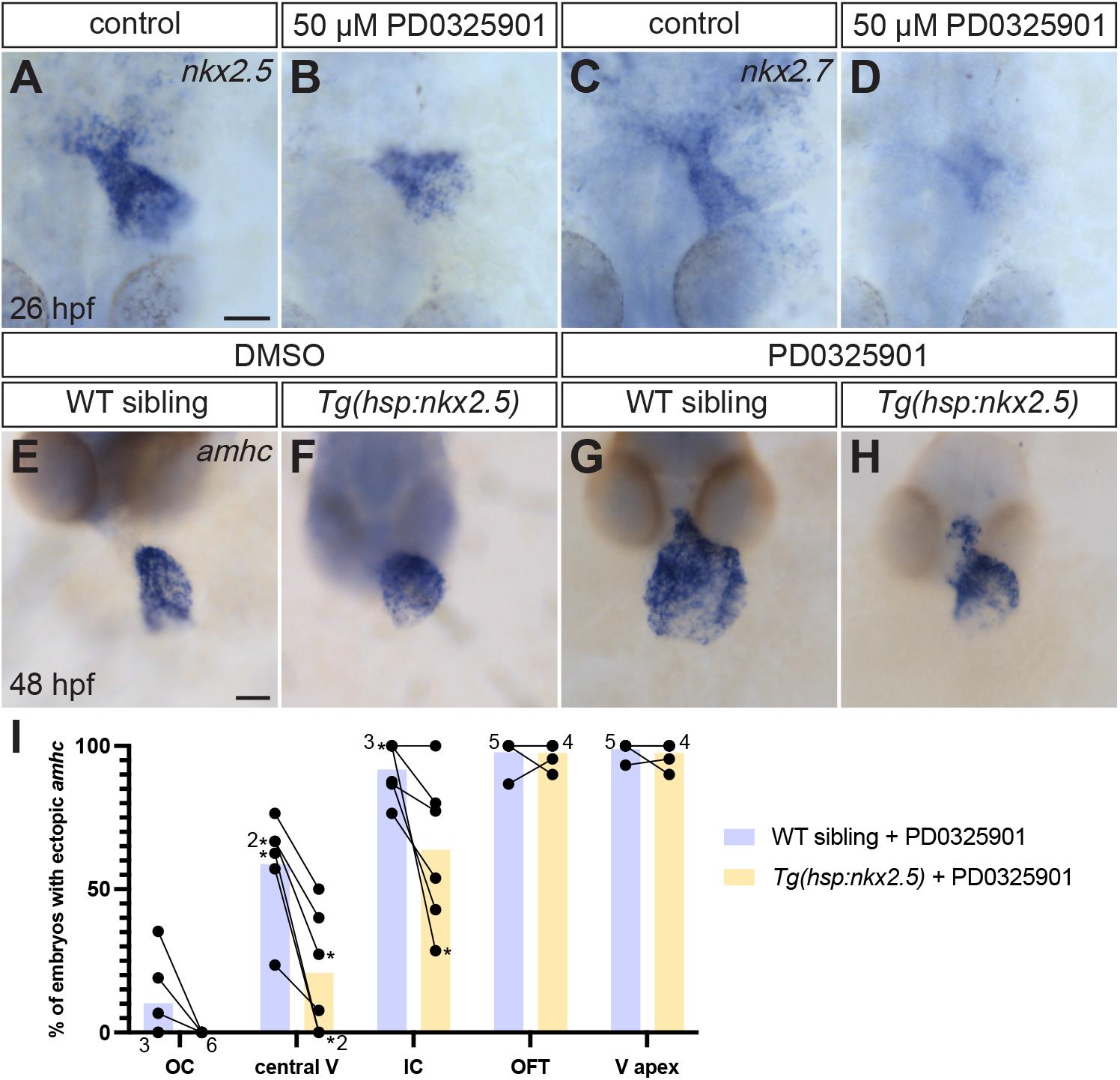
MEK-ERK signaling functions upstream of *nkx2.5* to maintain ventricular identity. (A-D) *In situ* hybridization shows expression of *nkx2.5* (A,B) or *nkx2.7* (C,D) at 26 hpf, dorsal views. Wild-type embryos were treated with DMSO or PD0325901 from 18 to 26 hpf. PD0325901-treated embryos (B,D) display reduced expression of *nkx2.5* and *nkx2.7* in the heart tube, compared to DMSO-treated controls (A,C). (E-H) Expression of *amhc* at 48 hpf (as in Fig. 1) indicates effects of heat shock at 17 hpf, followed by treatment with either DMSO (E,F) or 40-50 μM PD0325901 (G,H) from 18 to 22 hpf, as well as a subsequent heat shock at 22 hpf, in WT sibling (E,G) and *Tg(hsp:nkx2.5)* (F,H) embryos. (I) Graph (as in Fig. 5E) indicates percentage of PD0325901-treated embryos that expressed ectopic *amhc* in each ventricular region. Results of six independent experiments are shown. *p<0.05, Fisher’s exact test. (A,B) n=14, (C) n=15, (D,E) n=13, (F) n=19, (G,H) n(WT sibling, *Tg(hsp:nkx2.5)*)=(8,7), (17,13), (7,7), (15,22), (17,8), (21,10) in 6 independent experiments. Scale bars: 50 μm.

### Constitutive MEK1 activity induces ectopic *vmhc* expression in the atrium

While evaluating the impact of *Tg(hsp:caMEK1)* in the ventricle, we were intrigued to find that constitutive MEK1 activity also led to an interesting atrial phenotype. Specifically, we observed ectopic *vmhc* expression in the atrium of *Tg(hsp:caMEK1)* embryos (Fig. 7D). Individual *vmhc*-expressing cells, or small clusters of such cells, were scattered throughout the atrium, without any discernible regional pattern: ectopic *vmhc* expression was seen near the atrioventricular canal, near the inflow tract, or in the middle of the atrium, and each embryo exhibited a unique distribution of atrial *vmhc*. Ectopic *vmhc* expression emerged between 28 and 30 hpf (data not shown); by 30 hpf, multiple atrial *vmhc*-expressing cells were present (Fig. 7F) and appeared to be scattered in a manner similar to that seen at 48 hpf (Fig. 7D). Expression of *Tg(hsp:caMEK1)* robustly induced ectopic *vmhc* expression when heat shock was administered between 15 to 18 hpf, while the severity and penetrance of the phenotype was substantially reduced when embryos were heat shocked before 15 hpf or after 18 hpf (Fig. S6), demonstrating a discrete time window when constitutive MEK1 activity influences atrial expression of *vmhc*. Ectopic *vmhc* expression could also be induced by a second, independently derived allele of *Tg(hsp:caMEK1)*, indicating that this phenotype is caused by constitutive MEK1 activity and not by positional effects of the transgene (Fig. S7). Taken together, these data suggest that high levels of MEK-ERK signaling can disrupt the maintenance of atrial chamber characteristics.

**Figure 7.**
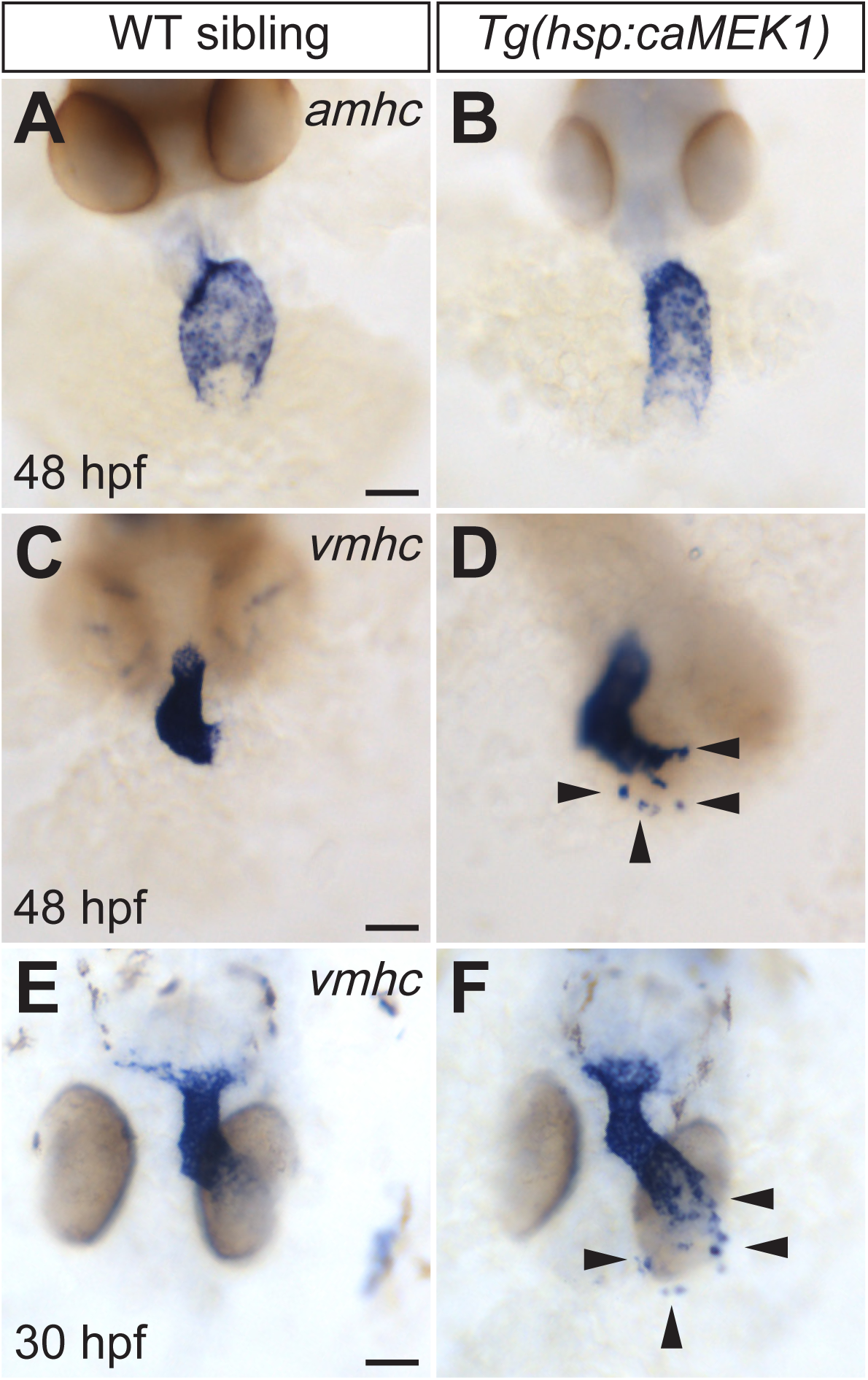
Constitutive MEK1 activity can induce ectopic *vmhc* expression. *In situ* hybridization shows expression of *amhc* (A,B) or *vmhc* (C-F) in frontal views at 48 hpf (A-D) or in dorsal views at 30 hpf (E,F), following heat shock at 16 hpf. In contrast to their WT siblings (C,E), most *Tg(hsp:caMEK1)* embryos exhibited ectopic *vmhc* expression in the atrium (D,F; arrowheads). (A) n=6, (B) n=10, (C) n=22, (D) n=15/21, (E) n=38, (F) n=22/32. Scale bars: 50 μm.

These results surprised us, since the induction of ectopic *vmhc* expression by constitutive MEK1 activity contrasts with our prior observation that increased FGF signaling, driven by expression of *Tg(hsp:cafgfr1)*, did not induce ectopic *vmhc* expression in the atrium (Pradhan et al., 2017). Revisiting and extending those experiments, we again failed to find ectopic expression of *vmhc* in *Tg(hsp:cafgfr1)* embryos after heat shock at 14, 16, or 18 hpf (Fig. S8 and data not shown). We wondered whether *Tg(hsp:caMEK1)* might be more effective than *Tg(hsp:cafgfr1)* at increasing MEK-ERK signaling, but we did not find evidence for this in our examination of pERK localization. Expression of *Tg(hsp:cafgfr1)* robustly increased the number of pERK+ cardiomyocytes and heightened the intensity of the pERK signal (Figs S9, S10), without any apparent deficiency compared to the effects of *Tg(hsp:caMEK1)* (Figs 4, S9, S10). However, we note that our analysis of pERK levels at a single timepoint would not reveal differences in the duration or dynamics of ERK activity, and we recognize that the kinetics of ERK activity are affected by multiple negative feedback mechanisms (Kim and Bar-Sagi, 2004; Lake et al., 2016; Szybowska et al., 2021; Thisse and Thisse, 2005). We expect that increased ERK signaling in response to *Tg(hsp:cafgfr1)* would be subject to negative feedback regulation by endogenous pathways that modulate MEK and ERK activity, including the FGFR endocytosis machinery and negative regulatory proteins such as Spry and Spred (Kim and Bar-Sagi, 2004; Lake et al., 2016; Szybowska et al., 2021; Thisse and Thisse, 2005), whereas ERK signaling in response to *Tg(hsp:caMEK1)* would not be attenuated by these factors. This presumed difference in susceptibility to negative feedback pathways could potentially account for the distinct impacts of *Tg(hsp:caMEK1)* and *Tg(hsp:cafgfr1)*.

We hypothesized that the ectopic *vmhc* expression observed in *Tg(hsp:caMEK1)* embryos represented atrial cardiomyocytes that failed to maintain their chamber identity. Consistent with this idea, we found that the ectopic *vmhc* expression colocalized with a myocardial marker (Fig. 8D-I), indicating that these *vmhc*-expressing cells resided within the atrial myocardium. Furthermore, we did not find evident loss of *amhc* expression in *Tg(hsp:caMEK1)* embryos (Fig. 7B), and the ectopic *vmhc*-expressing cells appeared to maintain normal levels of Amhc protein (Fig. 8M-R). These data, taken together with the scattered location of the ectopic *vmhc*-expressing cells, suggested that these cells were atrial cardiomyocytes in the process of acquiring ventricular characteristics, instead of ventricular cardiomyocytes that had migrated into the atrium. Thus, while endogenous levels of pERK are not sufficient to induce *vmhc* expression in the atrium, it seems that heightened levels of MEK-ERK signaling can override the mechanisms that normally enforce the maintenance of atrial chamber identity.

**Figure 8.**
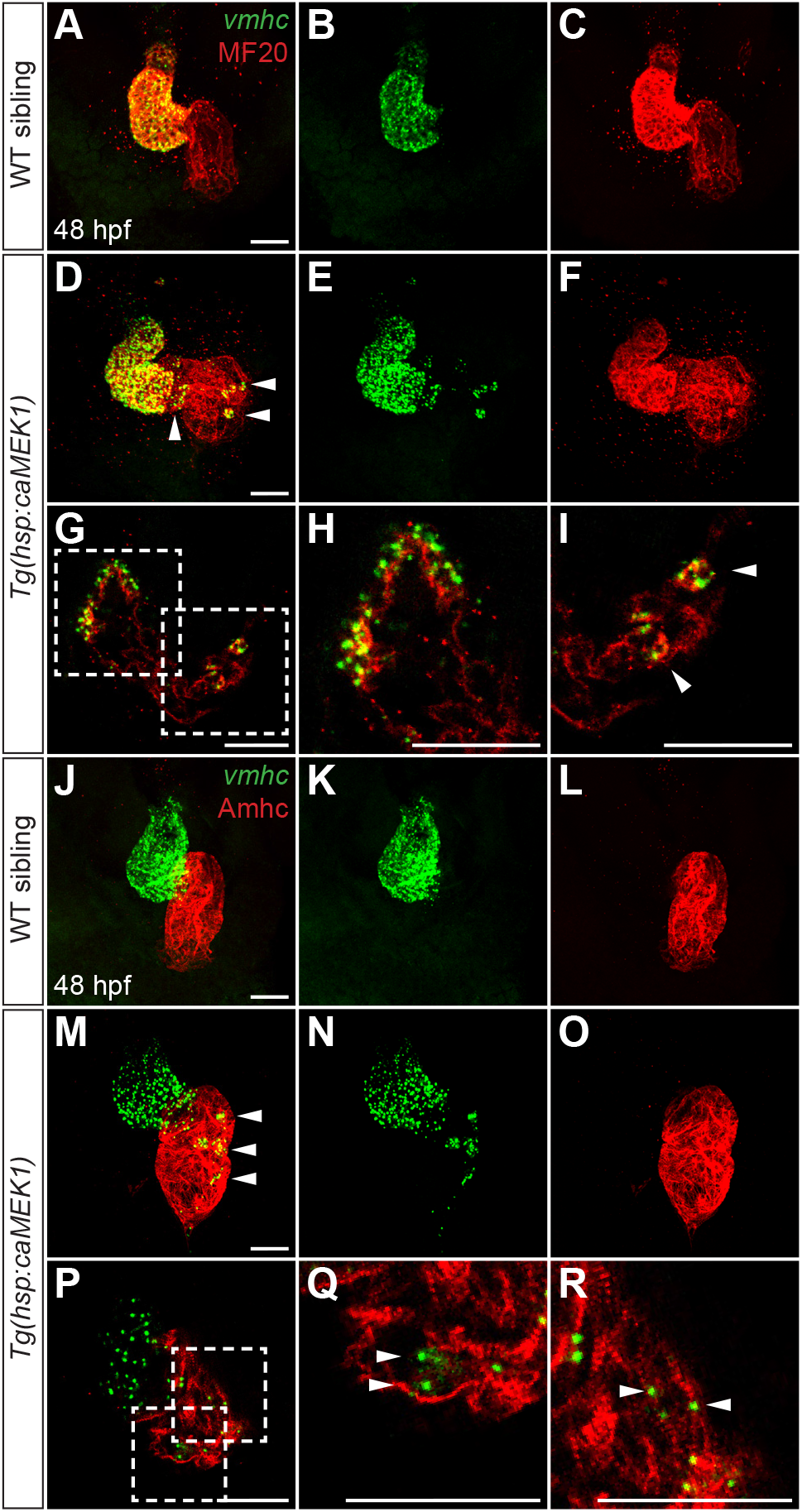
Constitutive MEK1 activity induces ectopic *vmhc* expression in the atrial myocardium. Fluorescent *in situ* hybridization shows expression of *vmhc* (green) together with immunofluorescence using MF20 (red, A-I) or labeling Amhc (red, J-R) at 48 hpf, following heat shock at 16 hpf. Three-dimensional reconstructions of frontal views (A-F, J-O), as well as single optical sections (G-I, P-R), highlight the localization of *vmhc* expression within the atrial myocardium in embryos expressing *Tg(hsp:caMEK1)* (D,E,G,I,M,N,P-R; arrowheads). Notably, in these experiments, all examined *Tg(hsp:caMEK1)* embryos exhibited ectopic *vmhc*, in contrast to the incomplete penetrance documented in Figure 7; this difference is likely due to the heightened sensitivity of fluorescent *in situ* hybridization. (A-C) n=6, (D-I) n=8/8, (J-L) n=5; (M-R) n=10/10. Scale bars: 50 μm.

## DISCUSSION

Our data indicate a previously unappreciated role for MEK-ERK signaling in enforcing the maintenance of ventricular chamber identity: MEK-ERK signaling is required after the onset of myocardial differentiation to repress ectopic induction of atrial genes and promote expression of ventricular genes in the developing ventricle, acting downstream of FGF signaling and upstream of *nkx2.5*. Intriguingly, our studies have also shown that the requirement for FGF-MEK-ERK signaling during chamber identity maintenance is context-dependent. Different portions of the ventricle exhibit differential degrees of sensitivity to inhibition of the FGF or MEK pathways, suggesting a regionalized pattern for ventricular plasticity. Meanwhile, in the atrium, we have uncovered a surprising ability of excessive MEK-ERK signaling to induce ventricular gene expression in atrial cardiomyocytes, even though endogenous levels of MEK-ERK signaling do not normally disrupt atrial identity. Taken together, our findings demonstrate distinct roles of MEK-ERK signaling in ventricular and atrial chamber identity maintenance.

Going forward, it will be important to identify additional components of the FGF-MEK-ERK-*nkx* pathway that promotes ventricular identity maintenance. It is not yet clear how directly ERK signaling impacts *nkx2.5* and *nkx2.7* expression or which effector genes might mediate this interaction. ETS transcription factors are interesting candidates for relevant effectors downstream of ERK signaling, since FGF signaling has been suggested to act through the Pea3 family of ETS factors in order to regulate the formation of *nkx2.5*-expressing cardiac progenitors in the early zebrafish embryo (Znosko et al., 2010). In addition to the identification of ERK-dependent regulators of *nkx* gene expression, it will be valuable to investigate whether there are other effector genes that act downstream of FGF-MEK-ERK signaling, in parallel with *nkx* genes, to regulate the maintenance of chamber-specific patterns of gene expression.

Analysis of additional players in the FGF-MEK-ERK-*nkx* pathway should also be coupled with further investigation of the dynamics of MEK-ERK signaling within the myocardium. The observed myocardial localization of pERK is consistent with a cell-autonomous role for MEK-ERK signaling in maintenance of ventricular cardiomyocyte characteristics, but the variable distribution and intensity of the pERK signal raise interesting questions about the duration and fluctuation of myocardial MEK-ERK signaling. In future studies, it will be beneficial to perform live imaging using ERK activity reporter transgenes, such as ERK-KTR sensors (De Simone et al., 2021; Qi et al., 2022; Simon et al., 2020), in order to evaluate precisely how myocardial MEK-ERK signaling is sustained or oscillates over time. These specific dynamics of MEK-ERK signaling may reveal regional patterns of signal intensity that were not apparent in our individual snapshots of pERK localization, and these patterns may be relevant to the differential plasticity of particular ventricular regions.

Furthermore, the regional differences in sensitivity to FGF-MEK-ERK pathway inhibition may reflect inherent distinctions between the differentiation states of specific portions of the ventricle. For instance, since OFT cardiomyocytes initiate their differentiation at a later stage than other groups of ventricular cardiomyocytes (de Pater et al., 2009; Hami et al., 2011; Lazic and Scott, 2011; Zhou et al., 2011), the late-differentiating OFT cells could be more vulnerable to modulation of the FGF-MEK-ERK pathway than the early-differentiating cells that are more advanced along their differentiation trajectory. This notion is consistent with our observation that inhibition of FGF or MEK signaling at 26 hpf still induces *amhc* expression in OFT cells without significantly impacting the OC, central ventricle, or IC (Fig. 1K,L). A similar logic could apply to the relative sensitivities of the OC and IC, since OC cardiomyocytes are thought to be more advanced in their differentiation than the relatively primitive cardiomyocytes in the IC (Auman et al., 2007; Christoffels et al., 2000; Habets et al., 2002; Jensen et al., 2013; Moorman and Christoffels, 2003) and may therefore require lower levels of FGF-MEK-ERK signaling to maintain their ventricular identity. In the future, it will be interesting to identify the differences in gene expression patterns or signaling pathway activities that distinguish portions of the ventricle and provide the molecular basis for regional differences in ventricular plasticity.

It is also intriguing to contemplate why the endogenous levels of pERK found in the atrial myocardium do not normally interfere with atrial identity maintenance, whereas heightened MEK signaling can induce ectopic expression of *vmhc* in the atrium. Perhaps there are atrium-specific repressive pathways that usually inhibit the effects of endogenous MEK-ERK signaling on chamber identity, such that only excessive levels of ERK signaling can override these repressive influences. Components of pathways that are known to regulate the maintenance of atrial identity, including the COUP-TFII, Nr2f1a, or TBX5 pathways (Martin et al., 2023; Sweat et al., 2023; Wu et al., 2013), might serve as such repressors. We note that the sparse expression of *vmhc* in the atrium of *Tg(hsp:caMEK1)* embryos contrasts with the broader expression of *amhc* in the ventricle of PD0325901-treated embryos. Interestingly, we did not observe a universal increase of pERK intensity in all of the cardiomyocytes in *Tg(hsp:caMEK1)* embryos. We therefore speculate that only the atrial cells with the most sustained or highest cumulative ERK activity over time went on to initiate *vmhc* expression in *Tg(hsp:caMEK1)* embryos; this idea seems consistent with the relatively small number of ectopic *vmhc*-expressing cells observed and the lack of a regional pattern for their location in the atrium. Ultimately, further experimentation will be needed to identify the chamber-specific signaling dynamics and molecular context that distinguish the role of MEK-ERK signaling in the atrium from its role in the ventricle.

Over the long term, we envision that strategies for cardiac tissue engineering will benefit from an enhanced understanding of how signaling pathways maintain cardiac chamber identities. There is ongoing interest in cardiac disease modeling and personalized drug discovery using engineered tissue derived from pluripotent cells (Huang et al., 2021; Lewis-Israeli et al., 2021; Parrotta et al., 2020; Protze et al., 2019). It is therefore critical to develop robust approaches for producing populations of ventricular and atrial cardiomyocytes that will stably maintain their chamber-specific characteristics (Ng et al., 2010; Zhao et al., 2020; Zhuang et al., 2022). While existing protocols are able to direct the initial differentiation of ventricular and atrial cells (Cyganek et al., 2018; Goldfracht et al., 2020; Lee et al., 2017; Zhao et al., 2019), the ongoing investigation of the mechanisms underlying chamber identity maintenance, including the roles of MEK-ERK signaling, will shape future approaches for continually enforcing the specific traits of ventricular and atrial tissue.

## MATERIALS AND METHODS

### Zebrafish

We used the following zebrafish strains: *Tg(myl7:EGFP) ^twu34^* (Huang et al., 2003), *Tg(kdrl:Hsa.HRAS-mCherry)^s896^* (Chi et al., 2008), *Tg(hsp70l:nkx2.5-EGFP)^fcu1^*(George et al., 2015), *Tg(hsp70:cafgfr1)^pd3^* (Marques et al., 2008). Embryos carrying *Tg(myl7:EGFP) ^twu34^* were identified by GFP fluorescence. Embryos carrying *Tg(kdrl:Hsa.HRAS-mCherry)^s896^*were identified by mCherry fluorescence. Embryos carrying *Tg(hsp70l:nkx2.5-EGFP)^fcu1^* were identified by GFP fluorescence following heat shock. Embryos carrying *Tg(hsp70:cafgfr1)^pd3^* were either identified by dsRed fluorescence in the lens at 48 hpf, or by PCR genotyping for *dsred* following immunofluorescence procedures, using the primers 5’-CAGTACGGCTCCAAGGTGTA-3’ and 5’-GTCCTCGAAGTTCATCACGC-3’, as in Figs S9 and S10. All zebrafish work followed protocols approved by the UCSD IACUC.

### Creation of stable transgenic lines

To generate transgenes for heat-activated expression of *MEK1^S219D^*, a constitutively active variant of zebrafish *map2k1* (ZDB-GENE-040426-2759), we first amplified the *MEK1^S219D^* coding sequence from a plasmid received as a gift from Dr. Chia-Hsiung Cheng (Chou et al., 2015) and cloned the amplicon into the middle donor vector pDONR221 (Invitrogen, 12536-017) through a Gateway BP reaction using the primers 5’- GGGGACAAGTTTGTACAAAAAAGCAGGCTACCATGCAGAAAAGGAGGAAGCC-3’ and 5’-GGGGACCACTTTGTACAAGAAAGCTGGGTGCATTCCCACACTGTGAGTCG- 3’. The transgene *Tg(hsp70:MEK1^S219D^-P2A-mCherry)* was then generated through a Gateway LR reaction (Invitrogen, 12537-023) containing four plasmids: pDONR221- MEK1^S219D^, p5E-hsp70l (Kwan et al., 2007), p3E-P2A-mCherry-pA (Covassin et al., 2009) and pDestTol2pA3 (Franco et al., 2019). We employed standard protocols to create transgenic founders (Soroldoni et al., 2009). Prospective founder fish were screened by evaluating mCherry fluorescence in their F1 progeny following heat shock, and phenotypic analysis was performed on the F2 and F3 progeny of two separate founders carrying distinct integrations of *Tg(hsp70:MEK1^S219D^-P2A-mCherry)*, both of which exhibited qualitatively similar levels of mCherry fluorescence. Similar phenotypes were observed in both transgenic lines. Data shown in Figs 4, 5, 7, 8, S6, and S10 depict results from the line *Tg(hsp70:MEK1^S219D^-P2A-mCherry)^sd70^*, and data shown in Fig. S7 depict results from the line *Tg(hsp70:MEK1^S219D^-P2A-mCherry)^sd71^*. In all experiments, embryos carrying *Tg(hsp70:MEK1^S219D^-P2A-mCherry)* were identified by mCherry fluorescence following heat shock.

### *In situ* hybridization

Standard whole-mount *in situ* hybridization and fluorescent *in situ* hybridization in combination with immunofluorescence were performed as previously described (Pradhan et al., 2017). We used established probes for the following genes: *amhc* (*myh6*; ZDB-GENE-031112-1), *vmhc* (*myh7*; ZDB-GENE-991123-5), *nkx2.5* (ZDB-GENE-980526-321) and *nkx2.7* (ZDB-GENE-990415-179). Location of ectopic *amhc* expression within the ventricle (Figs 2L, 5E,F, 6I,J) was categorized by a colleague who was blinded to the treatment condition or genotype of the samples.

### Immunofluorescence

Whole-mount immunofluorescence was generally performed as previously described (Pradhan et al., 2017), using antibodies listed in Table S1. For detection of pERK, we modified other established protocols (Lei et al., 2021; Watterston et al., 2021). Specifically, embryos were fixed in freshly made 4% PFA in PBS with PhosSTOP™(Roche, 4906845001) overnight at 4°C, then gradually dehydrated with a series of methanol/PBST washes before storage in 100% methanol at -20°C overnight. Embryos were then permeabilized in 50:50 acetone/methanol for 20 minutes at room temperature (RT) and re-hydrated through a series of methanol/PBST washes. Embryos were gradually warmed in 150 mM Tris-HCl (pH 9.0) buffer and were then kept at 70°C for 30 minutes. After cooling to RT, embryos were treated with Proteinase K (10 μg/ml) for 5 minutes (5 μg/ml if younger than 24 hpf). Embryos were next fixed in 4% PFA in PBS with PhosSTOP™ for 20 minutes at RT before being washed with PBST, blocked with dilution buffer (PBST with 2% BSA and 10% goat serum) for at least one hour at RT, and then incubated with primary antibodies for 2 nights at 4°C. Embryos were then washed with PBST, incubated with secondary antibodies overnight at 4°C in dark conditions, and washed in PBST at RT prior to imaging.

### Compound treatment

Compounds (see Table S2) were dissolved with 100% DMSO to a concentration of 10 mM and were stored in small aliquots at -20°C. Stock solutions were diluted to working concentrations (as indicated in figure legends) in E3 buffer. During treatment, embryos were incubated at 28.5°C in dark conditions. Control embryos were treated with E3 containing the same dilution of DMSO.

### Heat shock

To induce expression of *Tg(hsp70:MEK1^S219D^-P2A-mCherry)*, embryos were incubated in E3 buffer in a pre-warmed 37°C heat block for 1 hour. To induce expression of *Tg(hsp70l:nkx2.5-EGFP)*, embryos were incubated in E3 in a 37-37.5°C heat block for 30 minutes to an hour. To induce expression of *Tg(hsp70:cafgfr1)*, embryos were incubated in E3 in a pre-warmed 40°C incubator for 30 minutes, and they were then transferred to a pre-warmed 42°C heat block for 3 minutes.

### Imaging and image analysis

All bright-field images, as well as fluorescent images of embryos carrying *Tg(kdrl:Hsa.HRAS-mCherry)*, were captured using a Zeiss Axiocam and a Zeiss Axiozoom microscope and were processed using Zeiss AxioVision software. Confocal images were captured using a Leica SP5 or SP8 confocal laser-scanning microscope with a 25x water objective and a slice thickness of 1 µm, and confocal data were analyzed with Imaris software (Bitplane).

To evaluate pERK signal within the myocardium (Figs 3A’-D’, 4A’-D’, S3A’-D’, S4, S9A’-D’, S10), we created a ‘myocardial mask’ based on the intensity of either the *Tg(myl7:EGFP)* or CH1 signals, using the ‘surface’ function in Imaris. Only pERK signal within the boundary of the surface was extracted; however, this masking strategy might still include signals from tissues closely attached to the myocardium, such as the endocardium. Using the Imaris ‘spot’ function within the myocardial mask, we identified all of the cardiomyocytes in each sample and then counted the number of pERK+ cardiomyocytes that had an intensity mean value for the pERK channel above a certain threshold (Figs 3E, 4E, S9E). For each set of data acquired with the same settings, this threshold was set to include every spot that could be easily distinguished by eye from background fluorescence. We used a diameter of 5 μm when creating spots for all image analyses. To evaluate the intensity of the pERK signal (Figs 4F, S9F, S10), we first used the ‘spot’ function to mark all of the EGFP+ nuclei within the myocardial mask, since the EGFP generated by *Tg(myl7:EGFP)* expression typically accumulates in the nucleus at the stages examined. Then, to compare the overall pERK intensity between samples (Figs 4F, S9F), we obtained the mean of the intensity mean values for the pERK channel from all of the EGFP+ spots in each embryo. To examine the range of pERK intensity among cardiomyocytes (Fig. S10), we obtained intensity mean values for the pERK channel in each of the individual EGFP+ spots. Additionally, in order to compare the intensity of pERK signal between biological replicates (Figs 4F, S9F, S10), we normalized all of the values from wild-type samples and transgenic embryos such that the median of the wild-type values within each imaging session or clutch would be equal to 1 arbitrary unit (arb. unit).

### Statistics and replication

Statistical analysis was performed using Graphpad Prism 9. Non-parametric two-tailed Mann-Whitney tests were conducted when data involved a continuous variable (Figs 3I, 4E,F, S9E,F, S10). Fisher’s exact test was used when data involved categorical variables (Figs 5E, 6I). Asterisks in graphs are used to indicate statistical significance: * to indicate p<0.05, ** to indicate p<0.01, and **** to indicate p<0.0001. For *in situ* hybridization and immunofluorescence results without quantification, images shown are representative examples of multiple observations. The number of embryos examined, typically from at least two independent experiments, is specified in the figure legends.

## ACKNOWLEDGEMENTS

We thank C.H. Cheng for providing the *MEK1^S219D^*construct, H. Kwan for generating the *Tg(hsp70:MEK1^S219D^-P2A-mCherry)* plasmid, D. Gupta for performing blind analysis, and J. Diaz for creating the illustration in Fig. 2K. We also thank T. Sanchez, A. Yarbrough, G. Avillion, and the UCSD Animal Care Program for excellent zebrafish care. Finally, we thank K. Cooper, X. Sun, N.C. Chi, J. Chen, G. Guerrero, and the members of the Yelon lab for thoughtful input.

## COMPETING INTERESTS

No competing interests declared.

## AUTHOR CONTRIBUTIONS

Conceptualization: Y.Y., D.Y.; Methodology: Y.Y., D.Y.; Formal analysis: Y.Y., D.Y.; Investigation: Y.Y.; Writing - original draft: Y.Y., D.Y.; Writing - review & editing: Y.Y., D.Y.; Supervision: D.Y.; Project administration: D.Y.; Funding acquisition: Y.Y., D.Y.

## FUNDING

This work was supported by grants from the National Institutes of Health [R01 HL108599 and R01HL158877 to D.Y.], from the American Heart Association [21PRE827354 to Y.Y.], and from the UCSD Academic Senate [RG084137 to D.Y.].

## SUPPLEMENTARY FIGURES AND TABLES

**Supplementary Figure S1.**
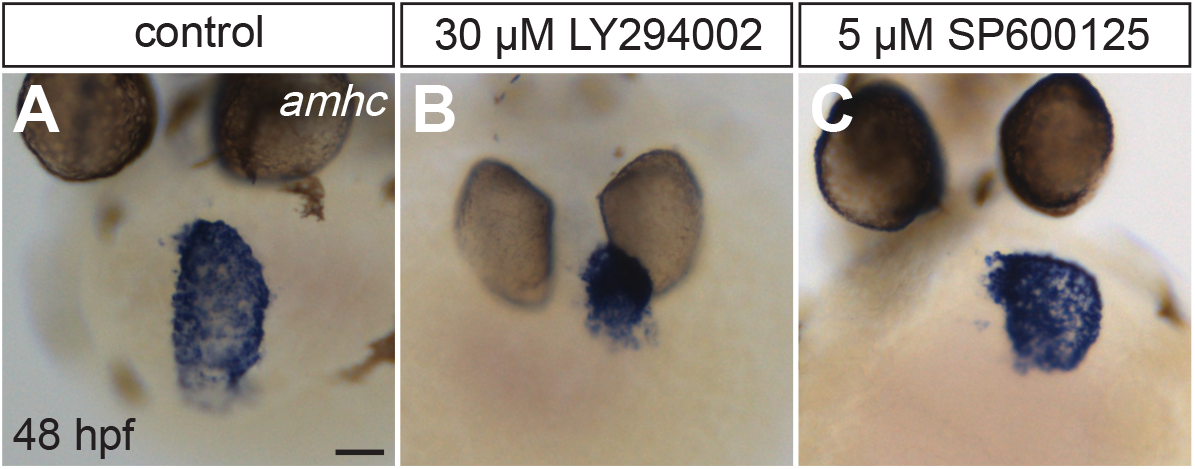
Inhibition of either the JNK pathway or the PI3K pathway fails to induce ectopic *amhc* in the ventricle. *In situ* hybridization shows expression of *amhc* at 48 hpf (as in Fig. 1) in wild-type embryos that were treated with DMSO (A), LY294002 (B) or SP600125 (C) from 18 (B) or 18.25 (C) to 48 hpf. (B) LY294002-treated embryos displayed a smaller atrium but no ectopic *amhc* in the ventricle. (C) SP600125-treated embryos did not display any ectopic *amhc* in the ventricle. Although neither inhibitor induced ectopic *amhc* expression, both were effective in causing previously reported morphological defects (Ma et al., 2009; Valesio et al., 2013) in treated embryos (data not shown). (A) n=9, (B) n=8, (C) n=13. Scale bar: 50 μm.

**Supplementary Figure S2.**
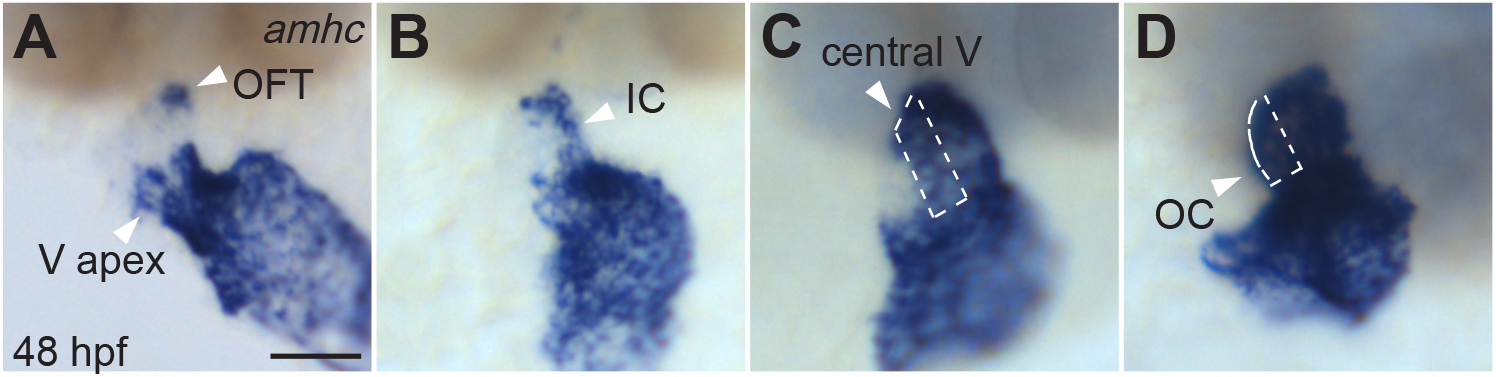
Ectopic *amhc* expression can be observed in different ventricular regions. *In situ* hybridization shows expression of *amhc* at 48 hpf (as in Fig. 1) in wild-type embryos that were treated with PD0325901 from 18 to 26 hpf. Arrowheads in (A) indicate examples of ectopic *amhc* expression in the ventricular apex (V apex) and outflow tract (OFT), and an arrowhead in (B) indicates an example of ectopic *amhc* expression in the inner curvature (IC). Areas marked by dashed outlines in (C) and (D) indicate examples of ectopic *amhc* expression in the central ventricle (central V) and outer curvature (OC).

**Supplementary Figure S3.**
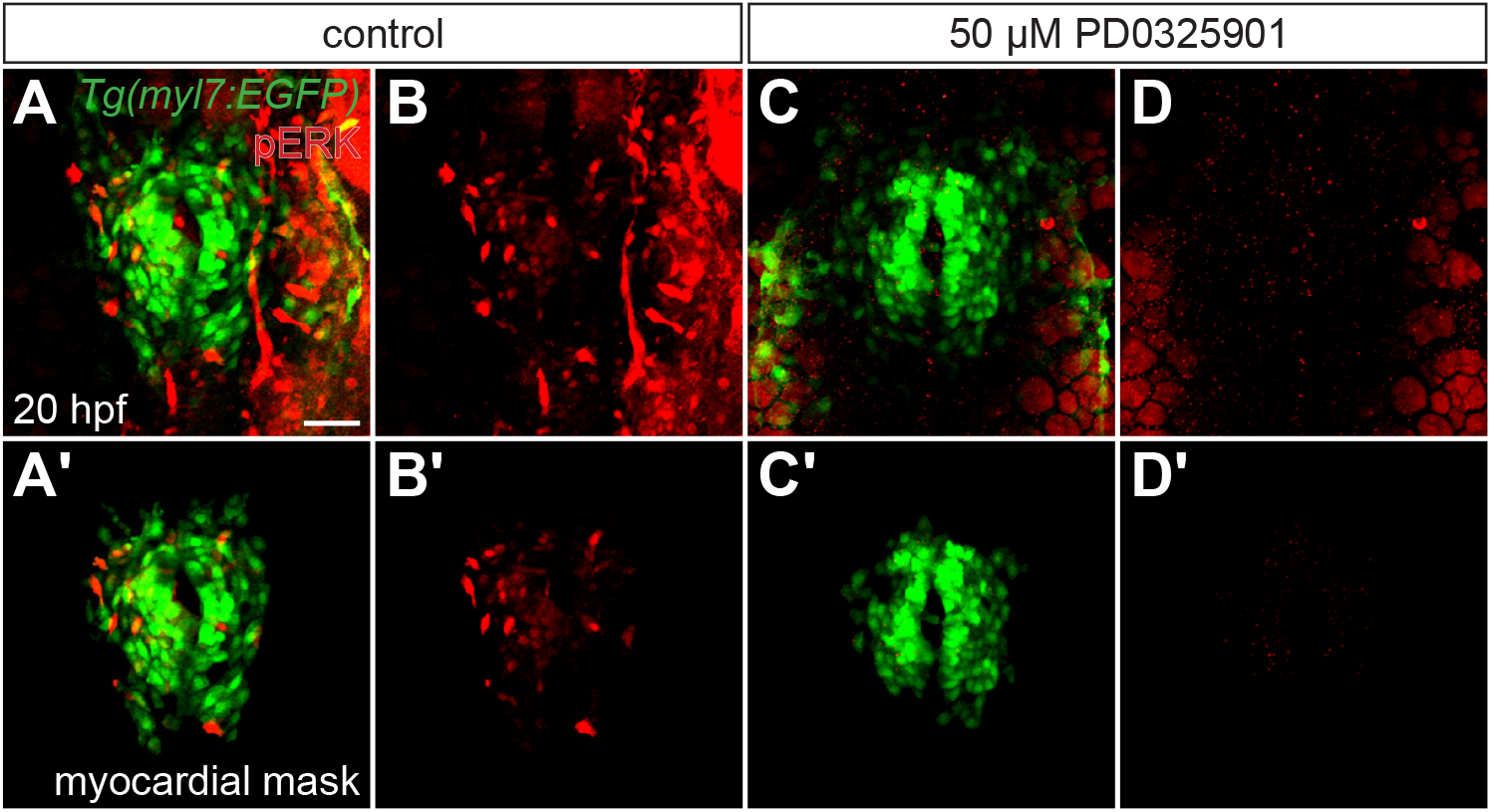
Treatment with PD0325901 effectively reduces pERK levels in the myocardium. Three-dimensional reconstructions compare pERK localization at 20 hpf (as in Fig. 3A,B) in *Tg(myl7:EGFP)* embryos treated with DMSO (A,B) or PD0325901 (C,D) from 16 to 20 hpf. Myocardial masks (A’-D’) are used as in Fig. 3A’,B’. Dorsal views, anterior up. PD0325901 treatment almost eliminates pERK signal (C,D), compared to what is observed in DMSO-treated embryos (A,B). (A-D) n=12. Scale bar: 50 μm.

**Supplementary Figure S4.**
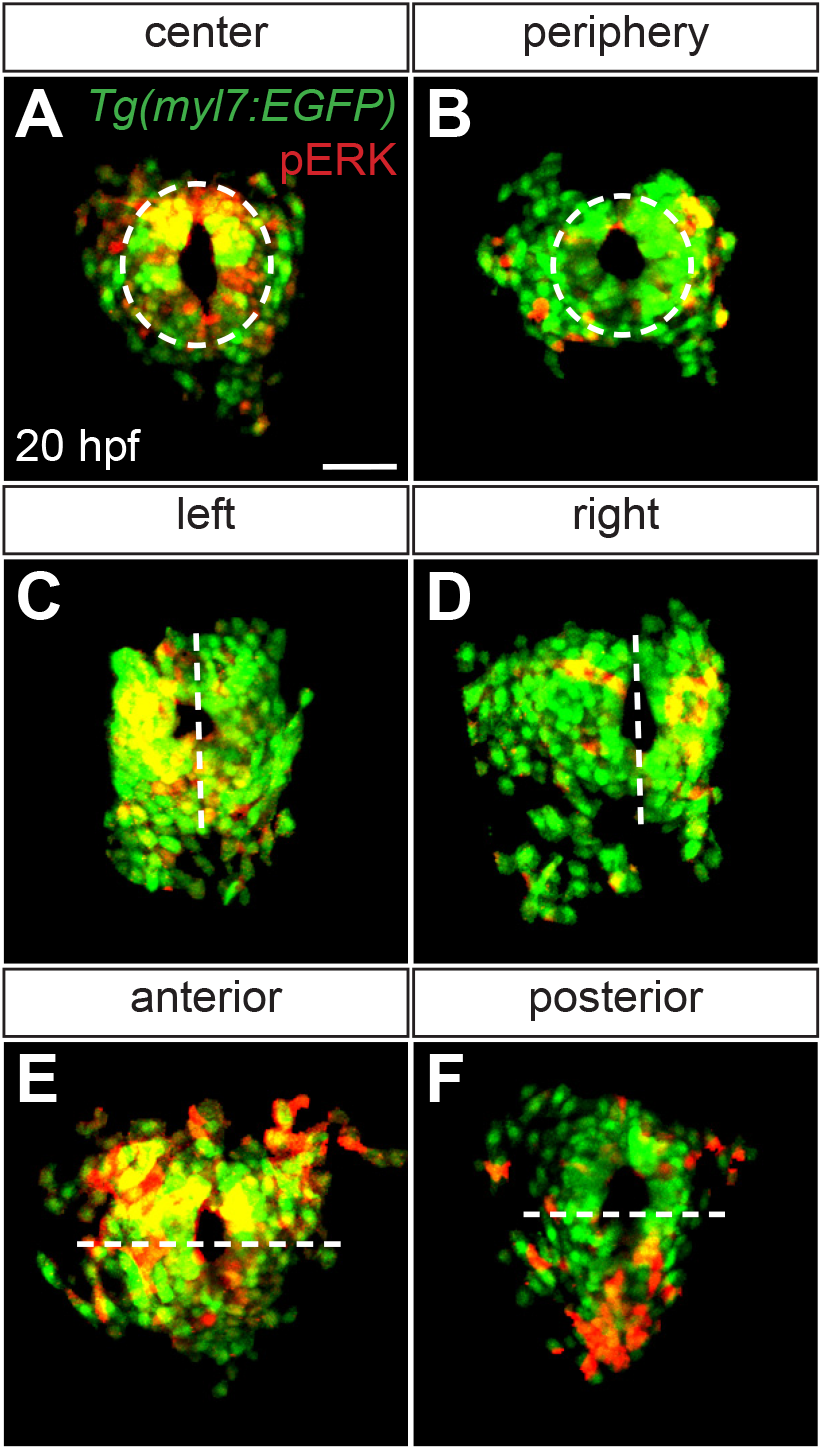
Variable distribution of pERK signal in the myocardium at 20 hpf. Additional examples of myocardial masks of three-dimensional reconstructions depicting pERK localization in *Tg(myl7:EGFP)* embryos at 20 hpf (as in Fig. 3 A’). Dorsal views, anterior up. In total, we have examined pERK localization in the myocardium of 75 wild-type embryos, and the distribution of pERK signal is unique in each. No regional patterns of myocardial pERK distribution are evident at 20 hpf. In the particular embryos shown here, pERK distribution appears enriched in the center (A) or along the periphery (B) of the myocardial cone, on the left (C) or right (D) side of the myocardium, in the anterior (E) or posterior (F) portion of the myocardium. Most embryos (79%) had comparable distribution of pERK in the center and in the periphery of the myocardial cone, whereas 4% of embryos exhibited enrichment in the center and 17% exhibited enrichment in the periphery. Many embryos (47%) had comparable distribution of pERK on the left and right sides of the myocardium, whereas 21% of embryos exhibited enrichment on the left and 32% exhibited enrichment on the right. Most embryos (68%) had comparable distribution of pERK in the anterior and posterior portions of the myocardium, whereas 20% of embryos exhibited enrichment in the anterior and 12% exhibited enrichment in the posterior. All assessments of enriched distribution were made qualitatively. Scale bar: 50 μm.

**Supplementary Figure S5.**
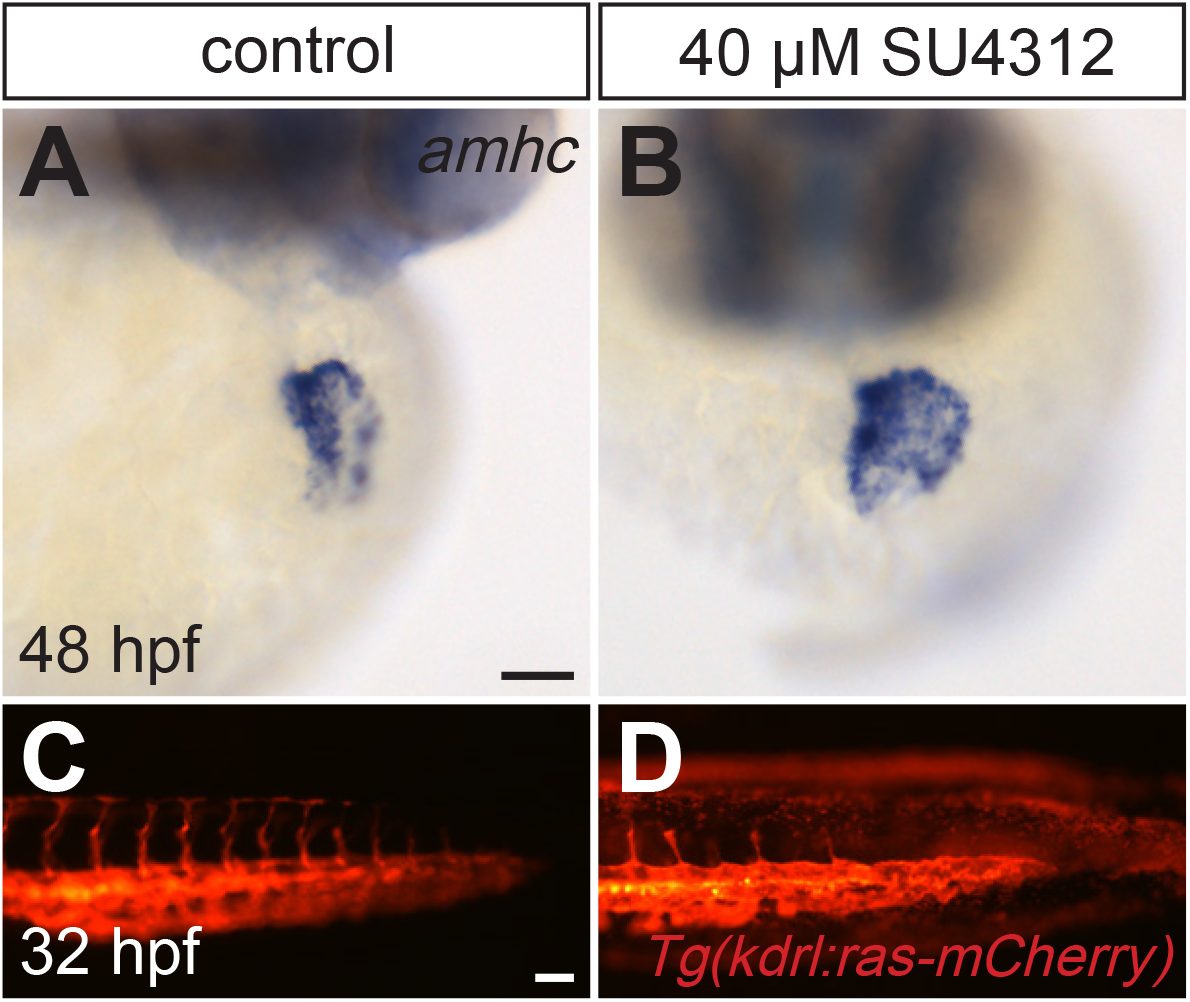
Inhibition of VEGF signaling does not affect maintenance of ventricular identity. (A,B) *In situ* hybridization shows expression of *amhc* at 48 hpf (as in Fig. 1) in embryos that were treated with SU4312 from 18 to 48 hpf. (C,D) Lateral views of the tail show expression of *Tg(kdrl:Hsa.HRAS-mCherry)* at 32 hpf in embryos that were treated with SU4312 from 24 to 32 hpf. SU4312-treated embryos did not exhibit ectopic *amhc* expression (B) but did exhibit impaired outgrowth of intersegmental vessels (D). (A) n=7, (B) n=11, (C, D) n=15. Scale bars: 50 μm.

**Supplementary Figure S6.**
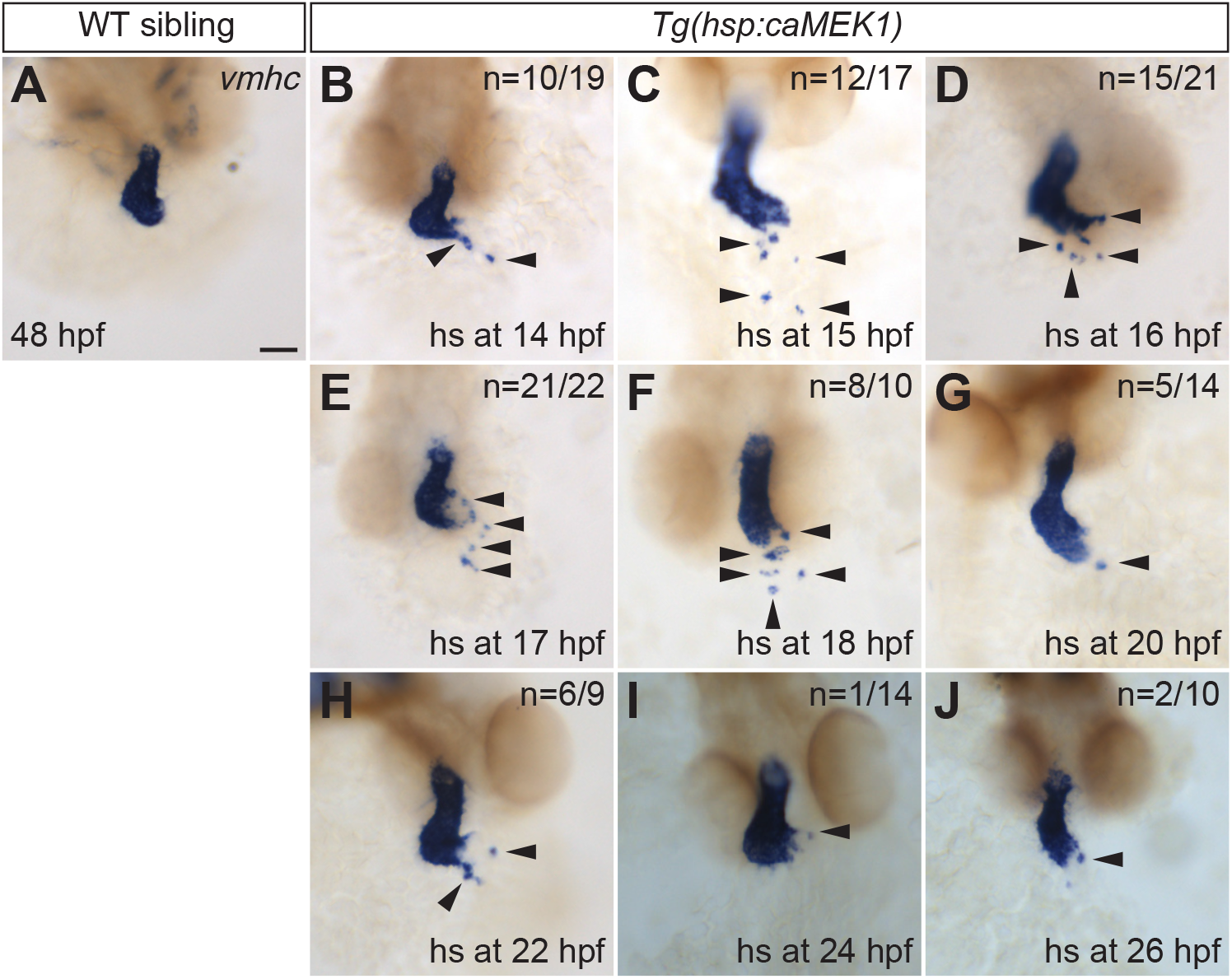
Constitutive MEK1 activity within a particular time interval induces ectopic *vmhc* expression. *In situ* hybridization shows expression of *vmhc* at 48 hpf (as in Fig. 1) in a WT sibling embryo (A) that was heat shocked at 14 hpf and in *Tg(hsp:caMEK1)* embryos (B-J) that were heat shocked at the indicated stages. For each stage of heat shock, the number of embryos exhibiting ectopic *vmhc* expression (arrowheads) is indicated. (A) n=20. Scale bar: 50 μm.

**Supplementary Figure S7.**
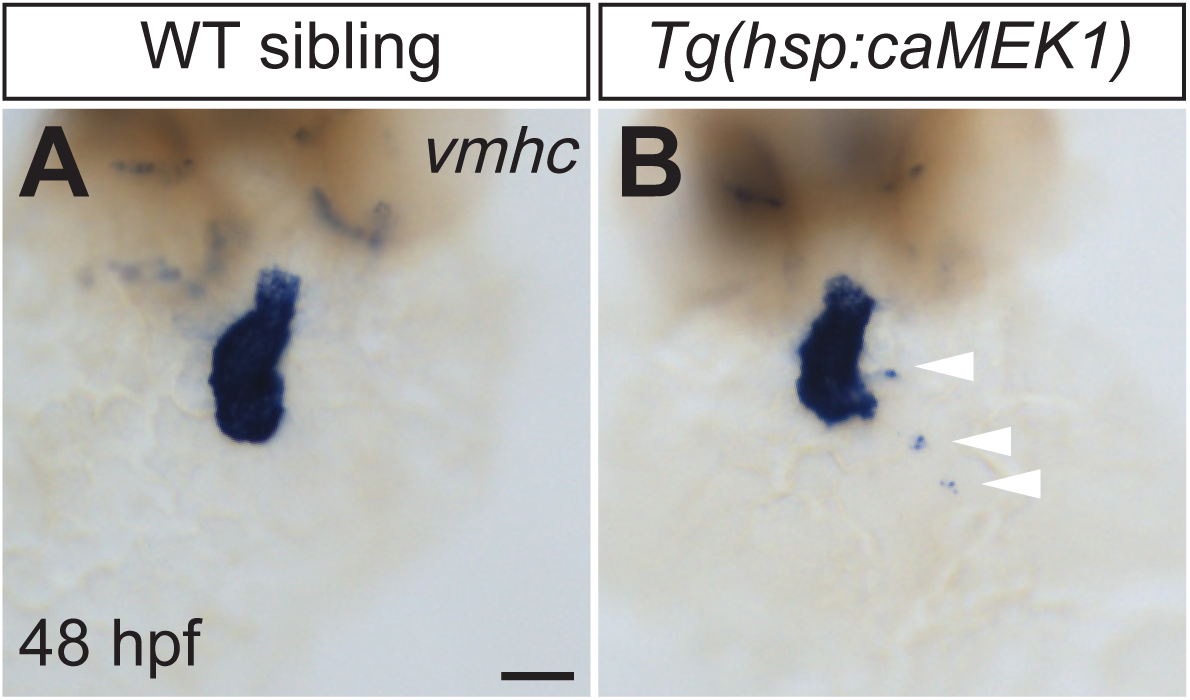
A second allele of *Tg(hsp:caMEK1)* demonstrates that constitutive MEK1 activity induces ectopic *vmhc* expression. (A,B) *In situ* hybridization shows expression of *vmhc* at 48 hpf (as in Fig. 1), following heat shock at 16 hpf. (B) *Tg(hsp:caMEK1)^sd71^* embryos exhibited ectopic *vmhc* expression (arrowheads). (A) n=25, (B) n=14/27. Scale bar: 50 μm.

**Supplementary Figure S8.**
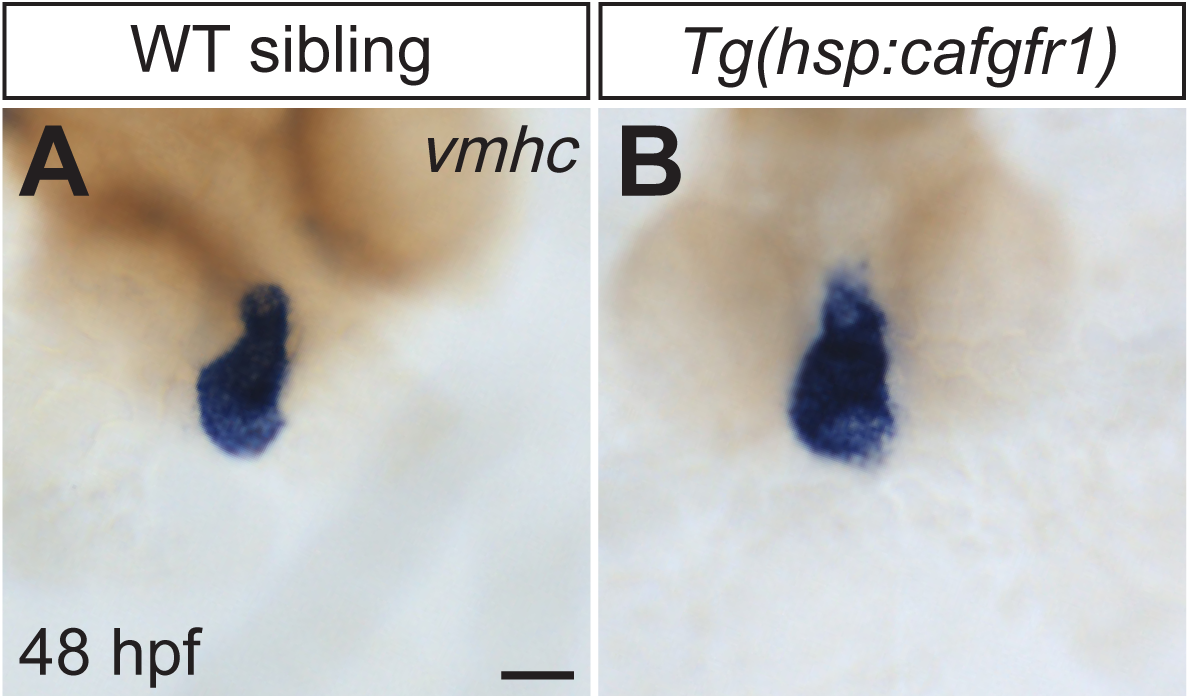
Constitutive activity of FGFR1 does not induce ectopic *vmhc* expression. (A,B) *In situ* hybridization shows expression of *vmhc* at 48 hpf (as in Fig. 1), following heat shock at 16 hpf. (B) *Tg(hsp:cafgfr1)* embryos did not exhibit ectopic *vmhc* expression. (A) n=9, (B) n=16. Scale bar: 50 μm.

**Supplementary Figure S9.**
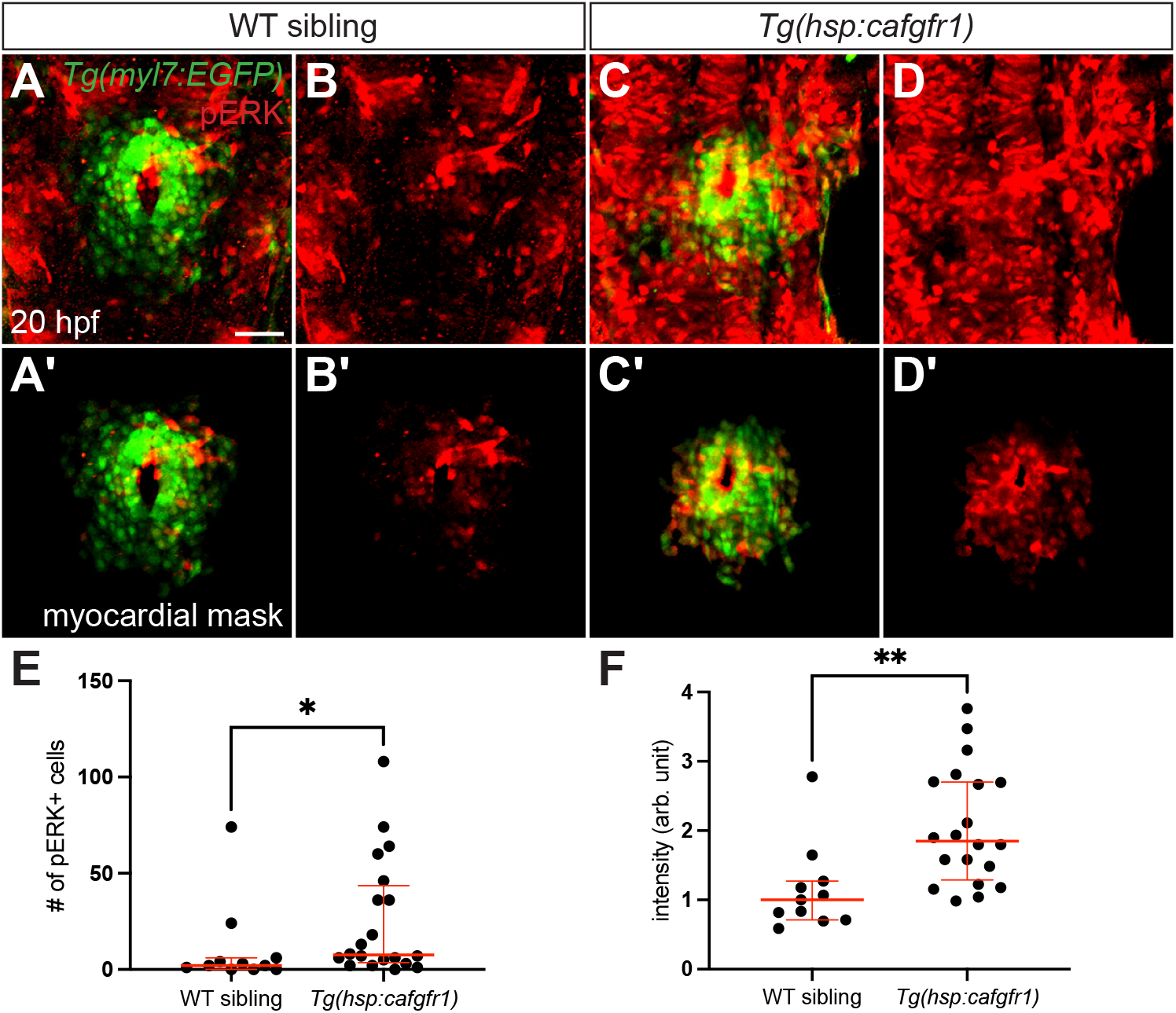
Constitutive FGFR1 activity increases pERK levels in the myocardium. Three-dimensional reconstructions compare pERK localization at 20 hpf (as in Fig. 4A-D) in *Tg(myl7:EGFP)* embryos (A,B) and *Tg(myl7:EGFP)* embryos carrying *Tg(hsp:cafgfr1)* (C,D), after heat shock at 16 hpf. Myocardial masks (A’-D’) are used as in Fig. 3A’,B’. Expression of *Tg(hsp:cafgfr1)* increases the amount of detectable pERK (C,D), compared to the endogenous level observed in WT sibling embryos (A,B). (A,B) n=11, (C,D) n=20. Scale bar: 50 μm. (E) Graph (as in Fig. 4E) compares the numbers of pERK+ cardiomyocytes in WT siblings and *Tg(hsp:cafgfr1)* embryos. (F) Graph (as in Fig. 4F) compares the average intensity of myocardial pERK signal in each WT sibling and *Tg(hsp:cafgfr1)* embryo examined. Red lines represent median and interquartile range. *p=0.0273 and **p=0.0014, two-tailed Mann-Whitney test.

**Supplementary Figure S10.**
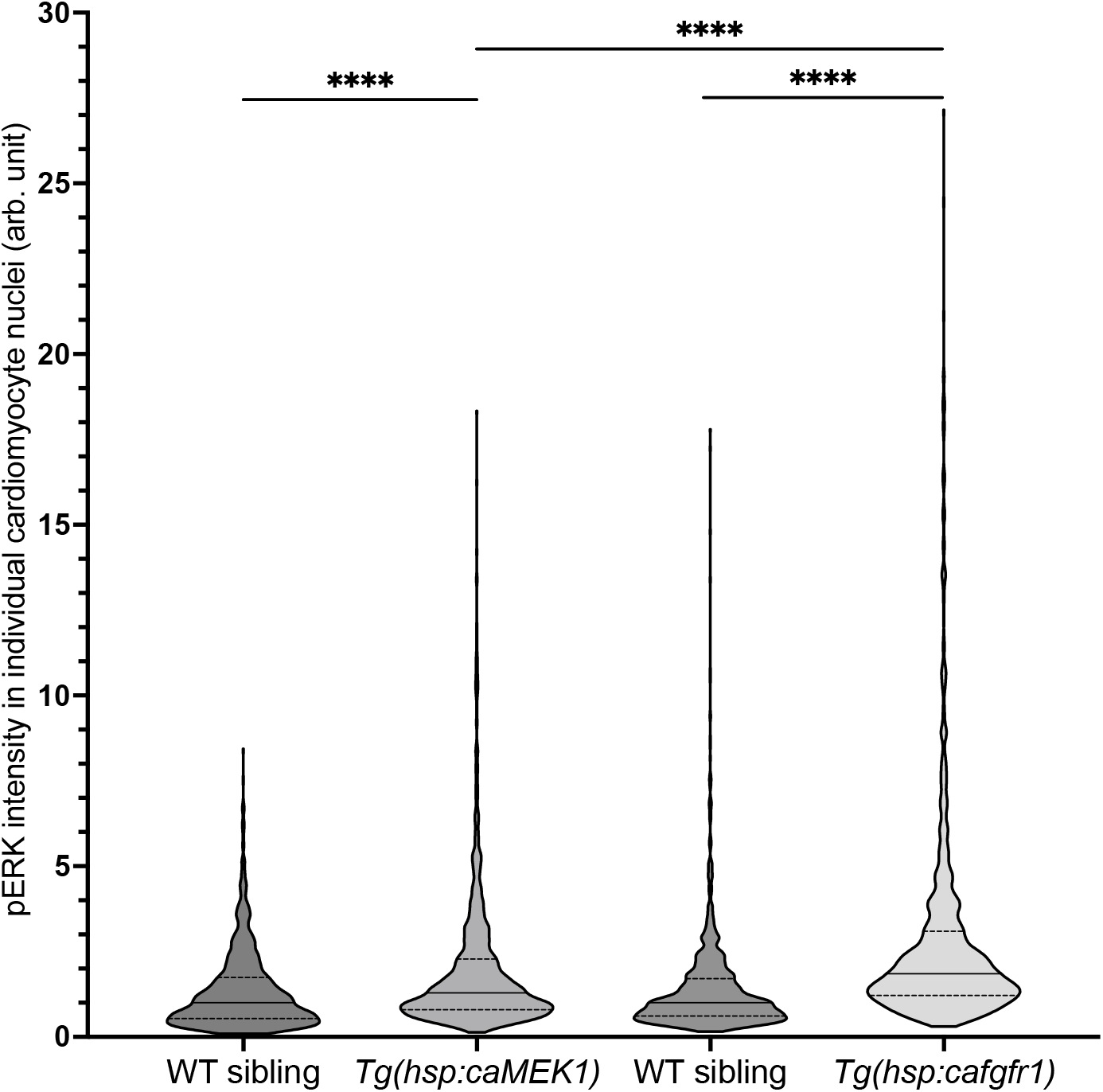
Constitutive activity of FGFR1 effectively increases pERK levels in the myocardium, without apparent deficiency compared to the effects of constitutive MEK1 activity. Violin plots compare the distribution of the intensity mean values of pERK signal from all of the cardiomyocyte nuclei examined in *Tg(hsp:caMEK1)* embryos, *Tg(hsp:cafgfr1)* embryos, and their corresponding WT siblings, all carrying *Tg(myl7:EGFP)*, at 20 hpf, following heat shock at 16 hpf. Normalized data are expressed in arbitrary (arb.) units (see Materials and Methods). Expression of *Tg(hsp:caMEK1)* or *Tg(hsp:cafgfr1)* shifted the distribution of pERK levels toward higher intensity, compared to the endogenous levels observed in WT sibling embryos, with a greater shift in *Tg(hsp:cafgfr1)* than in *Tg(hsp:caMEK1)*. In these experiments, laser power and exposure settings were even lower than those used for Figs 4 and S9, to facilitate capture of the full range of differences in pERK signals. Horizontal lines represent median and interquartile range. ****p<0.0001; two-tailed Mann-Whitney test. n=1263 nuclei in 11 *Tg(hsp:caMEK1)* embryos and 918 nuclei in 8 corresponding WT siblings, n=1145 nuclei in 10 *Tg(hsp:cafgfr1)* embryos and 1293 nuclei in 9 corresponding WT siblings.

**Supplementary Table S1.** Antibodies used for immunofluorescence.

Antibodies used for immunofluorescence
| Name | Source | Catalog # | Dilution |
| --- | --- | --- | --- |
| **Primary antibodies** |  |  |  |
| MF20 hybridoma supernatant | DSHB | MF20 | 1:10 |
| S46 hybridoma supernatant | DSHB | S46 | 1:10 |
| Phospho-Erk1/2 Rabbit mAb | Cell Signaling Technology | 4370 | 1:500 |
| Chicken anti-GFP IgY Ab | ThermoFisher Scientific | A10262 | 1:500 |
| CH1 hybridoma supernatant | DSHB | CH1 | 1:10 |
| **Secondary antibodies** |  |  |  |
| Goat anti-Mouse IgG (H+L) Ab, Alexa 594 | ThermoFisher Scientific | A11005 | 1:200 |
| Goat anti-Mouse IgG2b Ab, TRITC | Southern Biotech | 1090-03 | 1:100 |
| Goat anti-Mouse IgG1 Ab, FITC | Southern Biotech | 1070-02 | 1:100 |
| Goat anti-Rabbit IgG (H+L) Ab, Alexa 594 | ThermoFisher Scientific | A11012 | 1:500 |
| Goat anti-Rabbit IgG (H+L) Ab, Alexa 647 | ThermoFisher Scientific | A21245 | 1:500 |
| Goat anti-Chicken IgY (H+L) Ab, Alexa 488 | ThermoFisher Scientific | A11039 | 1:400 |
| Goat anti-Mouse IgG (H+L) Ab, Alexa 488 | ThermoFisher Scientific | A11029 | 1:500 |

**Supplementary Table S2.**
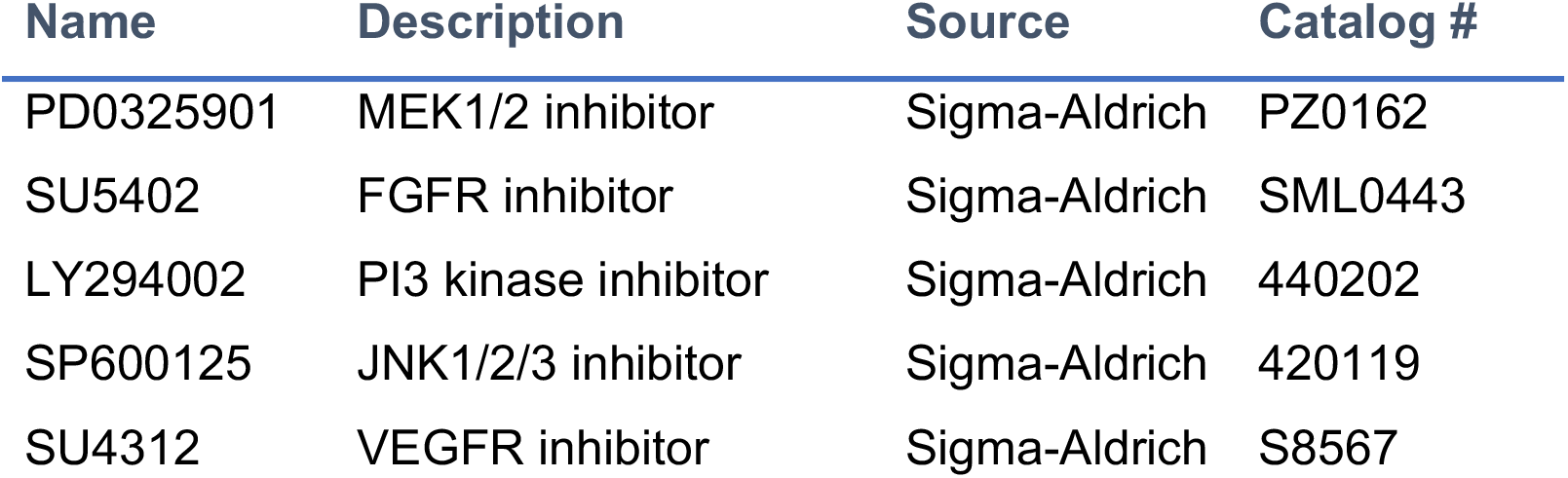
Compounds used to inhibit signaling pathways.

## Notes

### Competing Interest Statement

The authors have declared no competing interest.

